# Fitness consequences of structural variation inferred from a House Finch pangenome

**DOI:** 10.1101/2024.05.15.594184

**Authors:** Bohao Fang, Scott V. Edwards

## Abstract

Genomic structural variants (SVs) play a crucial role in adaptive evolution, yet their average fitness effects and characterization with pangenome tools are understudied in wild animal populations. We constructed a pangenome for House Finches, a model for studies of host-pathogen coevolution, using long-read sequence data on 16 individuals (32 *de novo-*assembled haplotypes) and one outgroup. We identified 643,207 SVs larger than 50 base pairs, mostly (60%) involving repetitive elements, with reduced SV diversity in the eastern US as a result of its introduction by humans. The distribution of fitness effects of genome-wide SVs was estimated using maximum likelihood approaches and showed SVs in both coding and non-coding regions to be on average more deleterious than smaller indels or single nucleotide polymorphisms. The reference-free pangenome facilitated discovery of a 10-million-year-old, 11-megabase-long pericentric inversion on chromosome 1. We found that the genotype frequencies of the inversion, estimated from 135 birds widely sampled geographically and temporally, increased steadily over the 25 years since House Finches were first exposed to the bacterial pathogen *Mycoplasma gallispecticum* and showed signatures of balancing selection, capturing genes related to immunity and telomerase activity. We also observed shorter telomeres in populations with a greater number of years exposure to *Mycoplasma*. Our study illustrates the utility of applying pangenome methods to wild animal populations, helps estimate fitness effects of genome-wide SVs, and advances our understanding of adaptive evolution through structural variation.

**Significance Statement:** Prevailing genomic research on adaptive and neutral evolution has focused primarily on single nucleotide polymorphisms (SNPs). However, structural variation (SV) plays a critical role in animal adaptive evolution, often directly underlying fitness-relevant traits, although their average effects on fitness are less well understood. Our study constructs a pangenome for the House Finch using long-read sequencing, capturing the full spectrum of genomic diversity without use of a reference genome. In addition to detecting over half a million SVs, we also document a large inversion that shows evidence of contributing to disease resistance. Our use of long-read sequencing and pangenomic approaches in a wild bird population presents a compelling approach to understanding the complexities of molecular ecology and adaptive evolution.

**Graphical abstract:** 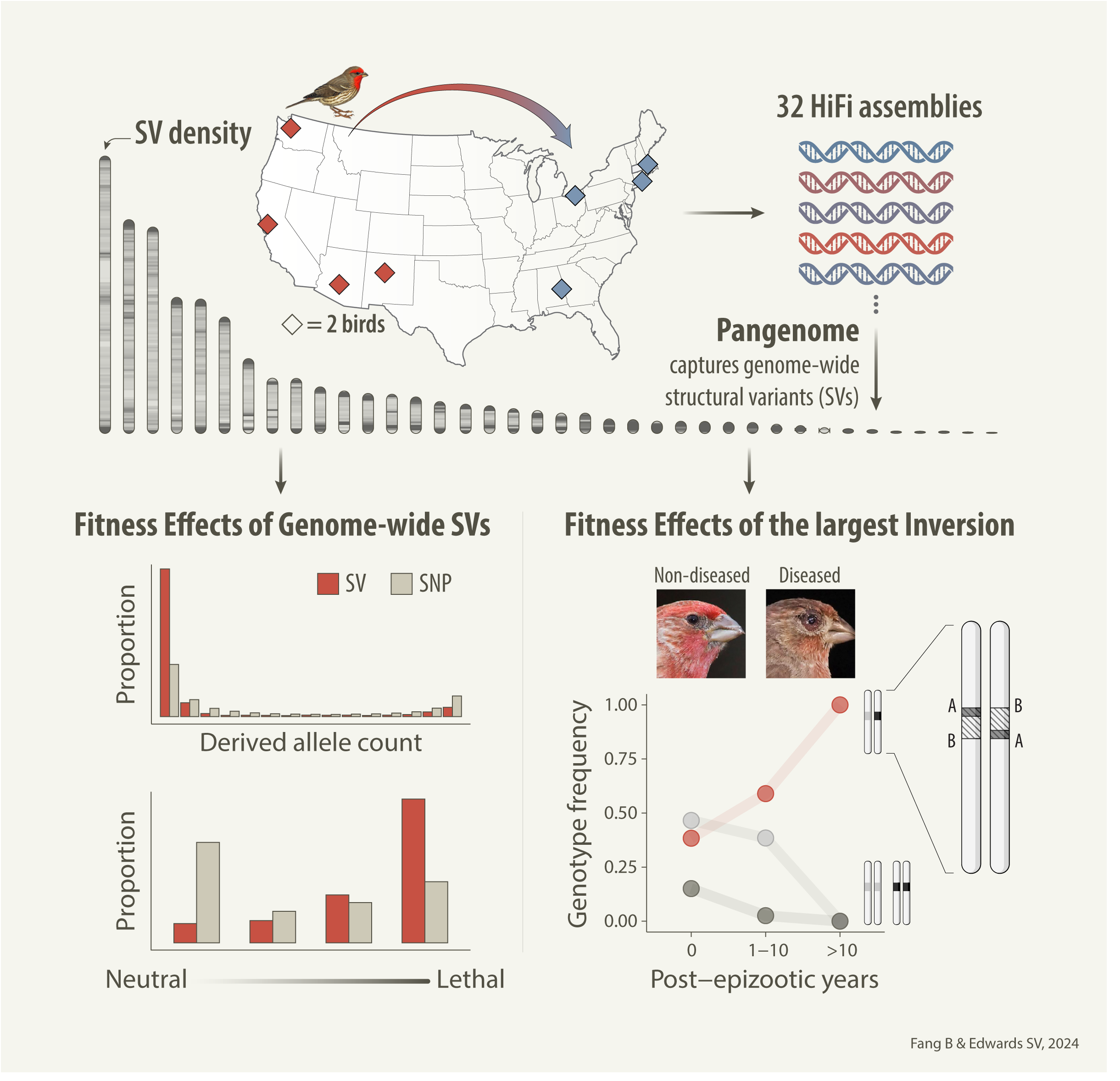

## Introduction

Structural variants (SVs) are large DNA changes consisting of insertions, deletions, and rearrangements in the genome. They differ from the more common and simpler ‘single nucleotide polymorphisms’ (SNPs), which involve only single DNA base changes. SVs are important for adaptation in natural populations (1–9), influencing morphology (10–13), behavior (14–17), and other physiological traits, such as disease resistance in humans (18), animals (8, 19), and plants (20). However, compared to SNPs, we know little about the fitness effects of SVs due to their underrepresentation in short-read sequencing data and the understudied links to phenotypes (4, 21–24).

One useful way of documenting SVs is the use of pangenomes. A pangenome models the complete set of genomic elements found within a species or clade, contrasting with reference-based methods, which compare sequences to a single genome, potentially biasing studies and missing variation due to choice of reference (25). Using long-read DNA sequencing methods, the totality of genetic variation in genomes can now be summarized in an organism’s pangenome, encompassing not only SNPs but also complex SVs (26, 27). Pangenome graphs, including de Bruijn and variation graphs, are considered superior data structures for categorizing extensive haplotypic genetic diversity within a single, data-rich model (25, 28–32). For instance, pangenome graphs have demonstrably improved mapping and discovery rates compared to reference genomes, allowing more comprehensive analysis of SVs (26, 33, 34). The application of pangenomic approaches is currently confined to humans (26, 34, 35), agricultural crops (36) livestock (37–41), and microbes (42, 43). Pangenomics in wild animals remains limited (44, 45), thus far mostly restricted by small sample sizes or focused on subspecies or higher taxonomic levels, making it difficult to evaluate the fitness effects of SVs.

The House Finch (*Haemorhous mexicanus*) is a model organism for studying host-pathogen coevolution; in the mid-1990s it experienced an epizootic involving a conjunctivitis-causing bacterial pathogen called *Mycoplasma gallisepticum* (MG) (46–51). House finches are native to the western US and were introduced to the eastern US by humans around 1940, where they adapted rapidly and began to expand west towards their original range (52). Following an MG outbreak in the eastern US from 1994–1998 and a subsequent outbreak in the western US in 2002, their resistance to the disease appears to have increased (53) as measured by gene expression (47, 54, 55), antibodies (56, 57), survival experiments (58, 59), and population surveys (53, 60). Here, we present a pangenome of the House Finch based on high-quality haplotypes assembled from 17 birds to characterize the full spectrum of genomic variants, including SVs, and investigate the fitness effects of SVs and their associations, if any, with adaptive disease resistance.

## Results

### House finch pangenome captures genome-wide structural variation

To construct the pangenome, we generated *de novo* assemblies for 16 House Finch samples and an outgroup, the Common Rosefinch (*Carpodacus erythrinus*; diverged 12.9 million years ago [Mya] (61, 62)), using PacBio HiFi long-read sequencing at approximately 42× coverage per bird (SI Appendix, Table S1, S2). We selected eight samples each from the western and eastern US (western and eastern, respectively) for a balanced genetic representation of the two populations (63) (Fig. 1B; Methods). The choice of the Common Rosefinch as the outgroup was driven by the limited availability of closely related species. Although closer outgroups such as the Purple Finch and Cassin’s Finch might be deemed appropriate, they represent a deep phylogenetic branch along with the House Finch within the Fringillidae family (64, 65), and might suffer from capturing variants still incompletely sorting between the outgroup and ingroup, which can complicate reconstruction of ancestral states. The somatic genome sizes of the House Finch and Rosefinch were similar (1.15 Gb and 1.19 Gb, respectively), suggesting that differences in genome size minimally impacted somatic variant discovery. We also incorporated a chromosome-level genome from a House Finch collected in California, produced and curated by the Vertebrate Genomes Project (VGP genome: 39 autosomes and sex chromosomes Z and W), primarily to provide a set of stable genomic coordinates in the pangenome and downstream analyses by us and other researchers. Our annotation strategies, which included in silico and evidence-based approaches (Methods), identified 22,080 protein-coding genes, 18.1% repeat content (SI Appendix Fig. S1) across the VGP genome. Two haplotypes per sample were assembled with hifiasm (66) (Fig. 1C; quality metrics in SI Appendix, Table S2). All data have been made publicly available (Data availability).

**Fig. 1.**
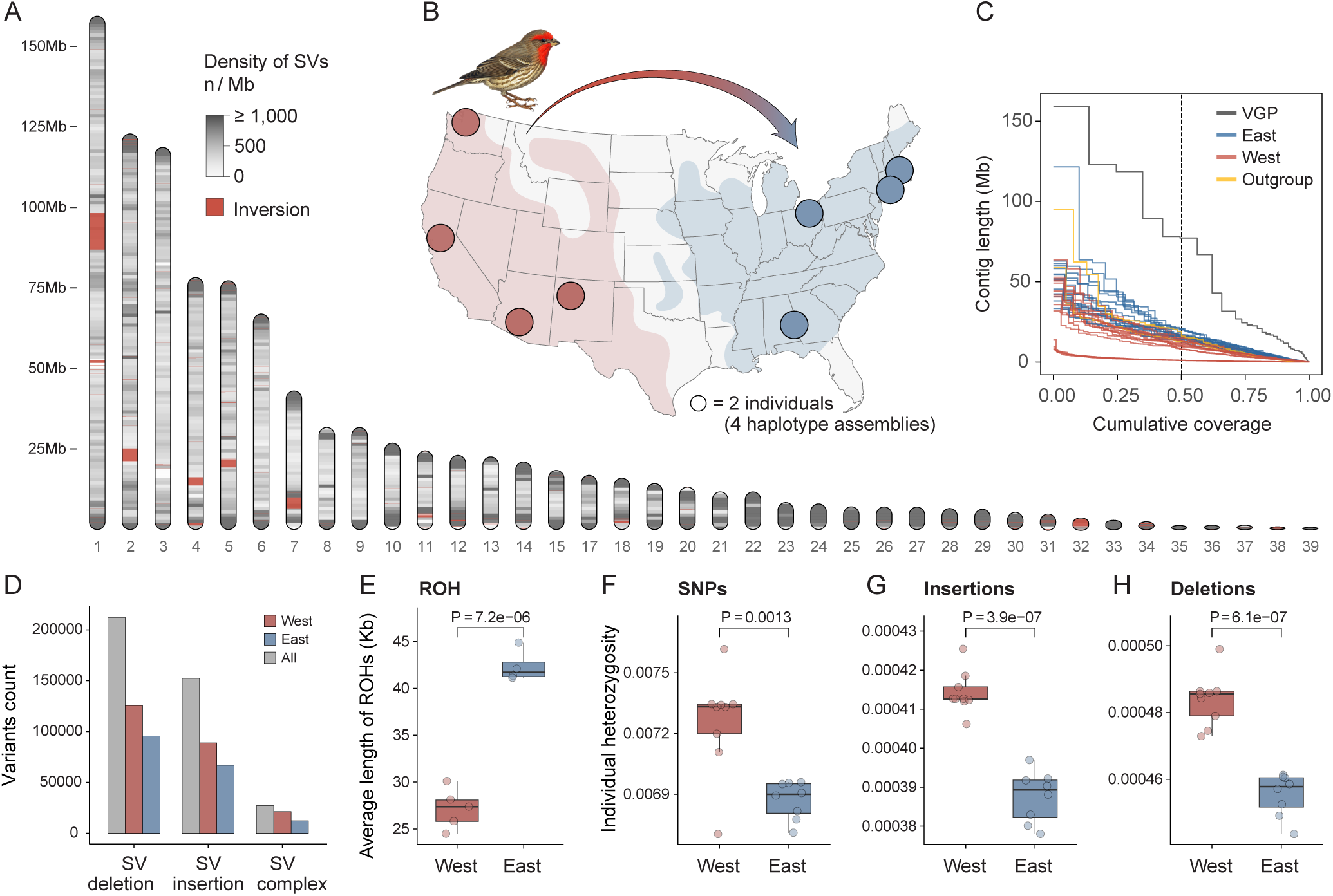
Genome-wide structural variants captured in the House Finch pangenome. (A) Genome-wide density of SVs larger than 50 bp. Sex chromosomes and chromosome 16 are not included in the analyses (see Methods). (B) Geographic distribution of 16 sampled House Finches. (C) Assembly contiguity of the 32 House Finch haplotype assemblies, a chromosome-level assembly from the Vertebrate Genomes Project (VGP), and two haplotype assemblies of the outgroup species Common Rosefinch (*Carpodacus erythrinus*), shown in a NGx graph. (D) Quantity of SVs in western and eastern House Finch populations extracted from the PGGB pangenome graph. (E) Population-wise runs of homozygosity (ROH). (F-H) Individual heterozygosity for SNPs (F), insertions (G), and deletions (H). *Bird illustration by Norman Arlott, © Lynx Edicions*.

The 35 haplotype assemblies, comprising 32 House Finch haplotypes, two Common Rosefinch haplotypes, and the VGP genome assembly, were used for pangenome construction. We built a separate pangenome graph for each the 38 autosomes using the PanGenome Graph Builder (PGGB) (30), a pipeline for constructing unbiased pangenome graphs using all-to-all genome alignments without relying on a reference genome. The selection of PGGB as the primary pangenome graph builder was based on its comprehensive ability to represent genomic variants of all sizes and its proven application in large-scale genomic projects (detailed justification in Methods). In the graph model, nodes represent DNA segments. Each node can orient in two ways: forward or reverse, creating a bidirected graph with four potential edges between any node pair to represent all combinations of orientations (SI Appendix Fig. S2). Haplotype sequences are depicted as paths within this graph. To characterize variants in the graph, we decomposed the graph to identify “bubble” subgraphs corresponding to non-overlapping variants (SI Appendix Fig. S2). In subsequent summaries of graph and variant statistics, we excluded the sex chromosomes and chromosome 16, which is comprised of 93% repeats (justification in Methods) and was fragmented even in the VGP assembly. Across 38 autosomes other than chromosome 16, the House Finch pangenome graph (including the outgroup) consists of 203,515,248 nodes, 286,630,623 edges, and 29,517 paths (see SI Appendix, Table S4 for graph statistics per chromosome). The pangenome coverage and growth curve indicate a plateau in the discovery of core variation shared by ≥90% of haplotypes, although the identification of unique variants continues to increase with the addition of more haplotypes (Fig. 2A).

**Fig. 2.**
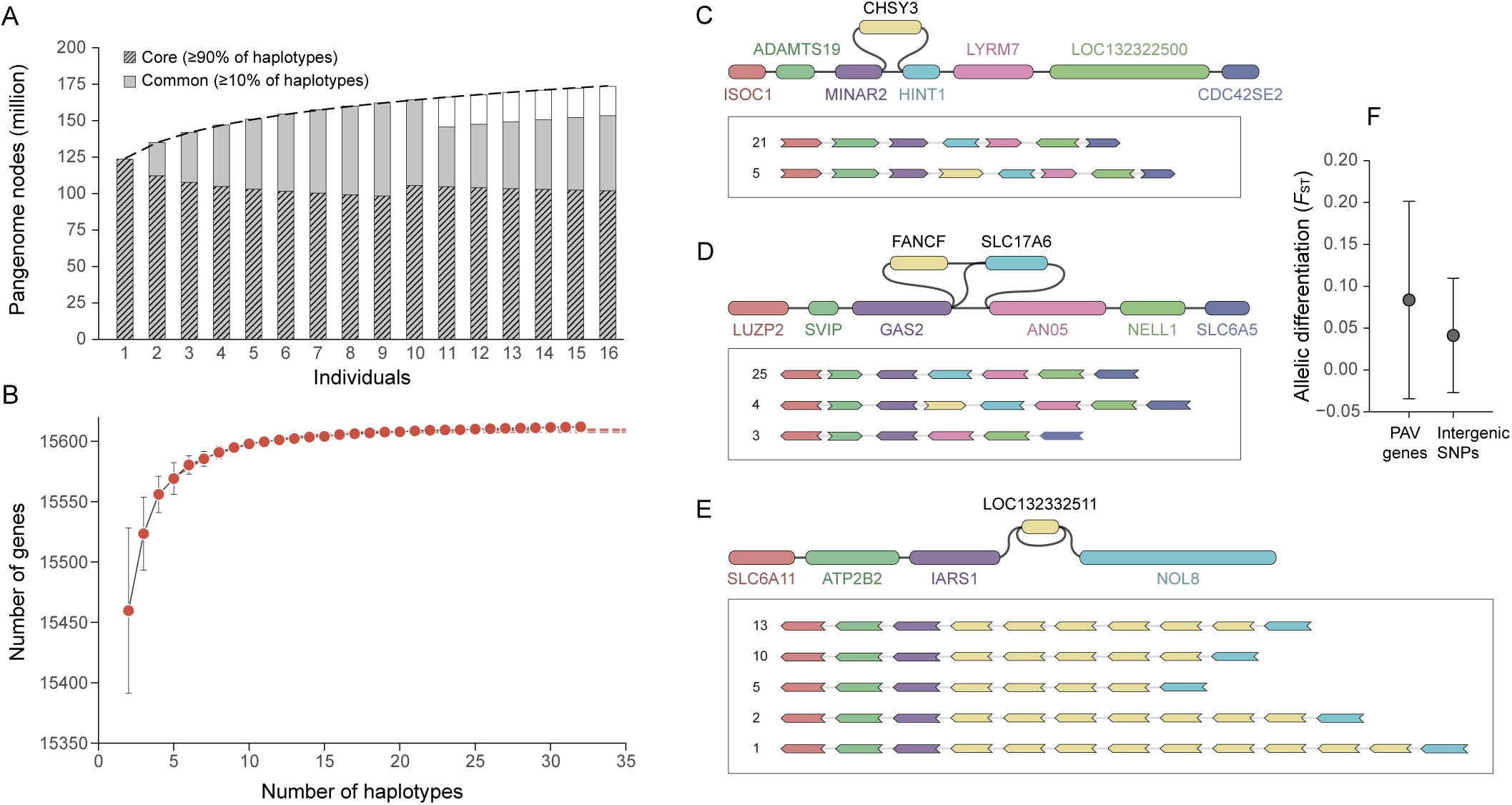
Pangenome and pangenome gene graph variation. (A) PGGB pangenome growth curves show number of new nodes (Y-axis) each sample (X-axis) adds to the graph. Nodes are classified as “core” if present in ≥90% of samples and “common” if present in ≥10%. (B) Evaluation of the pangenome gene plateau based on pangene. Data points indicate the average gene count across sampled haplotypes with 100 permutations. The red curve, extrapolating from observed data via a logistic growth model, forecasts the trend. The number of genes obtained from 25, 30, and 32 haplotypes suggest convergence and a plateau in the discovery of new genes. Genes from 32 haplotypes are annotated with miniprot (Methods). (C–E) Examples of gene presence–absence variation (PAV; C–D) and copy number variation (CNV; E) within the pangenome gene graph constructed using Pangene (Methods). In the graph, a node represents a gene and an edge between two genes indicates their genomic adjacency on the haplotypes. Each panel shows a subgraph of the variant gene, adapted from Bandage visualization (top), and distinct haplotypes (paths) within the subgraph visualized using pangene (bottom). Pangene filters out fragmented contigs. (F) Allele differentiation (*F*_ST_; mean ± S.D) of gene presence-absence polymorphisms and intergenic SNPs between western and eastern populations.

Decomposing the PGGB pangenome graph, we classified variants into 20,590,976 SNPs across House Finch haplotypes (Methods); 3,416,519 INDELs (<50 base pairs [bp]) including 1,308,590 insertions (INS) relative to the outgroup, 2,048,950 deletions (DEL), 58,979 complex INDELs (INDEL-complex); and 391,717 SVs (≥50 bp; Fig1D), including 152,234 SV-insertions (SV-INS), 212,256 SV-deletions (SV-DEL), and 27,227 complex SVs (SV-complex). Complex variants are defined as sites with different allelic sizes that are not 1 bp in length and/or not polarized by the outgroup (Methods). Multiallelic variants (824,858 SNPs, 725,479 INDELs and 251,490 SVs) were also identified (SI Appendix, Table S5). To enhance discovery of inversions and other SVs, and to compare the results of different methods for detecting SVs, we additionally used minigraph (28), a pangenome graph builder tailored to identify large SVs, as well as SVIM-asm (67) and SyRi (68), the latter two being reference-based SV callers (Methods; SI Appendix, Table S6).

Using these tools we identified 343 inversions ranging in size from 50 bp to 11.3 megabases (Mb) (Fig. 1A; SI Appendix, Table S6), with 163 inversions (48%) identified by at least two programs (SI Appendix Fig. S3). We manually confirmed all six large (>1 Mb) inversions using dot plots in HiFi contigs (Methods; examples in SI Appendix Fig. S4). We also identified 4,518 segmental duplications, ranging in size from 1,000 to 4,786,886 bp, using BISER (69), accounting for 8% of the VGP genome (SI Appendix, Table S3).

The SVs have a mean size of 174 bp and a median size of 338 bp, and encompass 6.5 times more base pairs than do SNPs across the genome (386,804,299 vs. 59,310,087 bp). Most SVs reside in introns and intergenic regions (SI Appendix Fig. S5). The largest SV was a 11.3 Mb inversion located in chromosome 1 (Chr1: 86,909,536 – 98,195,717), with the putative centromere located inside the inversion, thus constituting a pericentric inversion (Methods). Over half of the SVs (60%) span repeats as identified by RepeatMasker (70) (Methods), predominantly simple repeats (16.9%), long terminal repeats (LTRs, 3.2%), long interspersed nuclear elements (LINEs, 2.0%), and unclassified repeats (28.1%), with LTRs being particularly enriched in SVs of around 640 bp (Fig. 3B-D). Nearly 8% (7.7%) SVs span more than one repeat type (Fig. 3B).

**Fig. 3.**
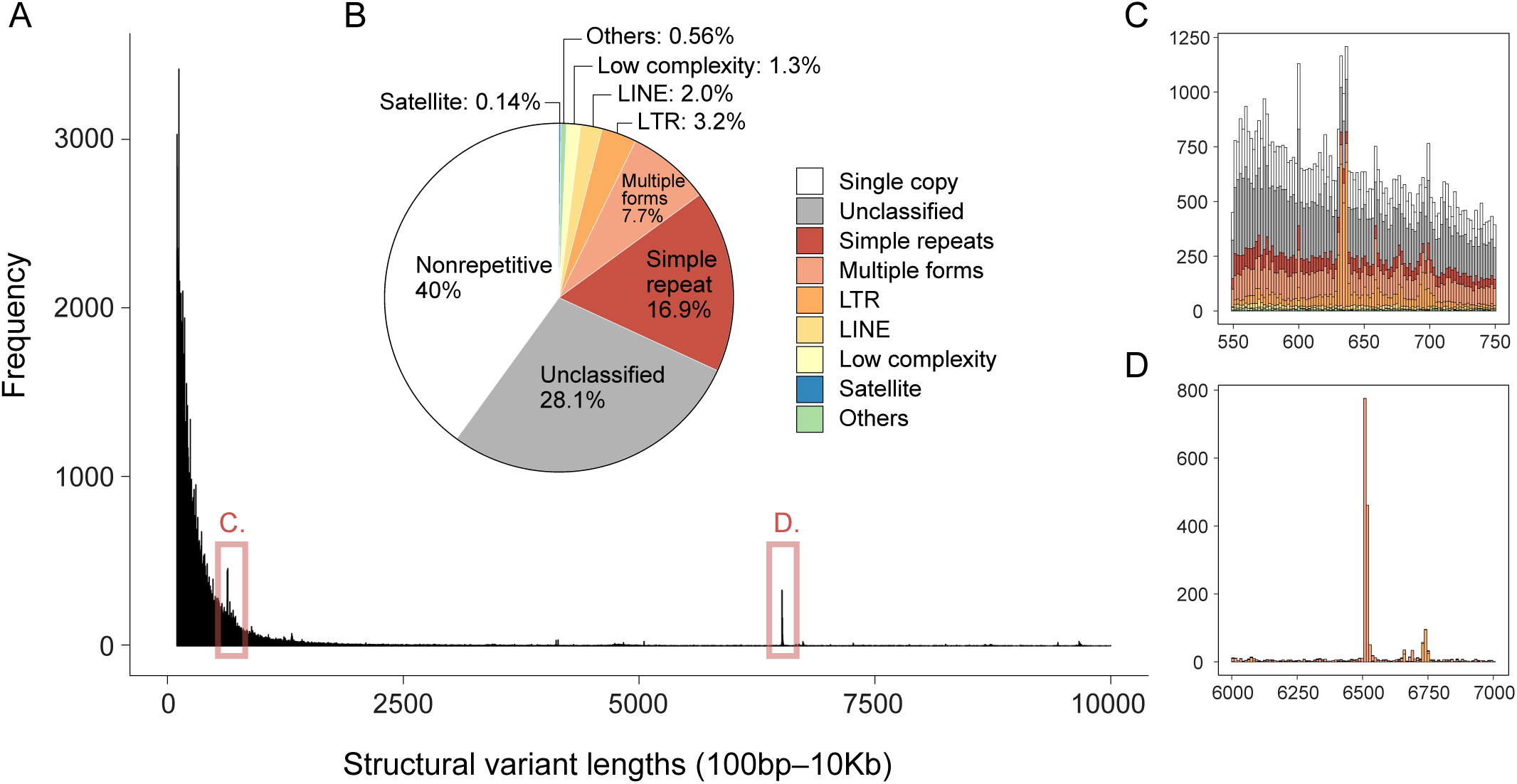
Most structural variants overlap repeats and transposable elements. (A) Histogram illustrating the distribution of lengths of structural variants (SVs), from 100 base pairs (bp) to 10 kilobases (Kb). (B) Pie chart delineating SV overlap with various repeat types, transposable elements and non-repeats. (C, D) Detail of histogram sections from panel A, displaying SV frequency in the 550–750 bp range (C) and the 6–7 Kb range (D), with color-coding corresponding to the keys for repeat classification.

### Pangenome gene graph captures genome-wide variation in genic copy number

To examine genic variation, including genic presence-absence variants (PAV) and copy number variation (CNV), we used Pangene (71) and an annotation of protein-coding genes for the 32 House Finch haplotypes to produce a pangenome gene graph (GFA format) in which each node represents a gene, and an edge between two genes indicates their genomic adjacency on the haplotypes. The annotations were based on the VGP genome using miniprot (72), which aligns proteins to each haplotype assembly. To reduce false positive gene variation caused by fragmented HiFi contigs, which Pangene filters out and thus flags as potential gene absence, and due to the challenges of mapping protein sequences to genomes (72, 73), we considered confident CNV genes as those with more than two copies in a haplotype, and confident PAV genes as absent in at least 5 haplotypes (85% filtering threshold, SI Appendix Fig. S6). The resulting graph includes 711 PAV genes, representing 4.5% of the genes annotated by miniprot (Fig. 2C, D), and 180 CNV genes, constituting 1.16% of the annotated genes (Fig. 2E). Allelic differentiation (*F*_ST_) of PAV genic polymorphisms between the eastern and western populations is low, although slightly higher than that for intergenic SNPs (Fig. 2F). However, outliers (*F*_ST_ > 0.4) were manually inspected as potential false positives due to fragmented contigs (SI Appendix Fig. S7). These suggest that the distribution of PAV genes may be driven largely by genetic drift.

### Reduced SV and SNP diversity in eastern birds

The human-mediated introduction of the House Finch to the eastern is known to have facilitated a reduction in genetic diversity compared to the ancestral western population (51, 74). Our data confirms evidence of reduced SNP diversity and also reveals decreased diversity SVs in the eastern population. This is evident in the fewer variants of all types found in the eastern population, with more SNPs, INDELs, and SVs found exclusively in the west than in the east (Fig. 1D). Additionally, the western population exhibits higher heterozygosity and lower levels of inbreeding, as indicated by shorter runs of homozygosity (ROH; Fig. 1E), than the eastern population. To compare individual heterozygosity between the two populations, we identified genomic regions deemed safe for calling SNPs and INDELs by Dipcall (75) (Methods). The results show significantly higher heterozygosity in the western population, measured across SNPs, insertions, and deletions (Fig. 1F-H).

### Fitness effects of genome-wide structural variants

We assessed the impact of genome-wide variants on fitness by analyzing their distribution of fitness effects (DFE). We focused on SNPs, INDELs, and SVs in multiple genomic contexts including coding (CDS), non-coding (intron and intergenic), and regulatory (untranslated region, 5’ UTR and 3’ UTR) regions. We estimated the DFE based on the site frequency spectrum (SFS; Fig. 4A), accounting for neutral demographic effects, using two maximum likelihood approaches, fastDFE (76) and anavar (77), with anavar providing enhanced correction for polarization errors and fastDFE offering greater computational efficiency.

**Fig. 4.**
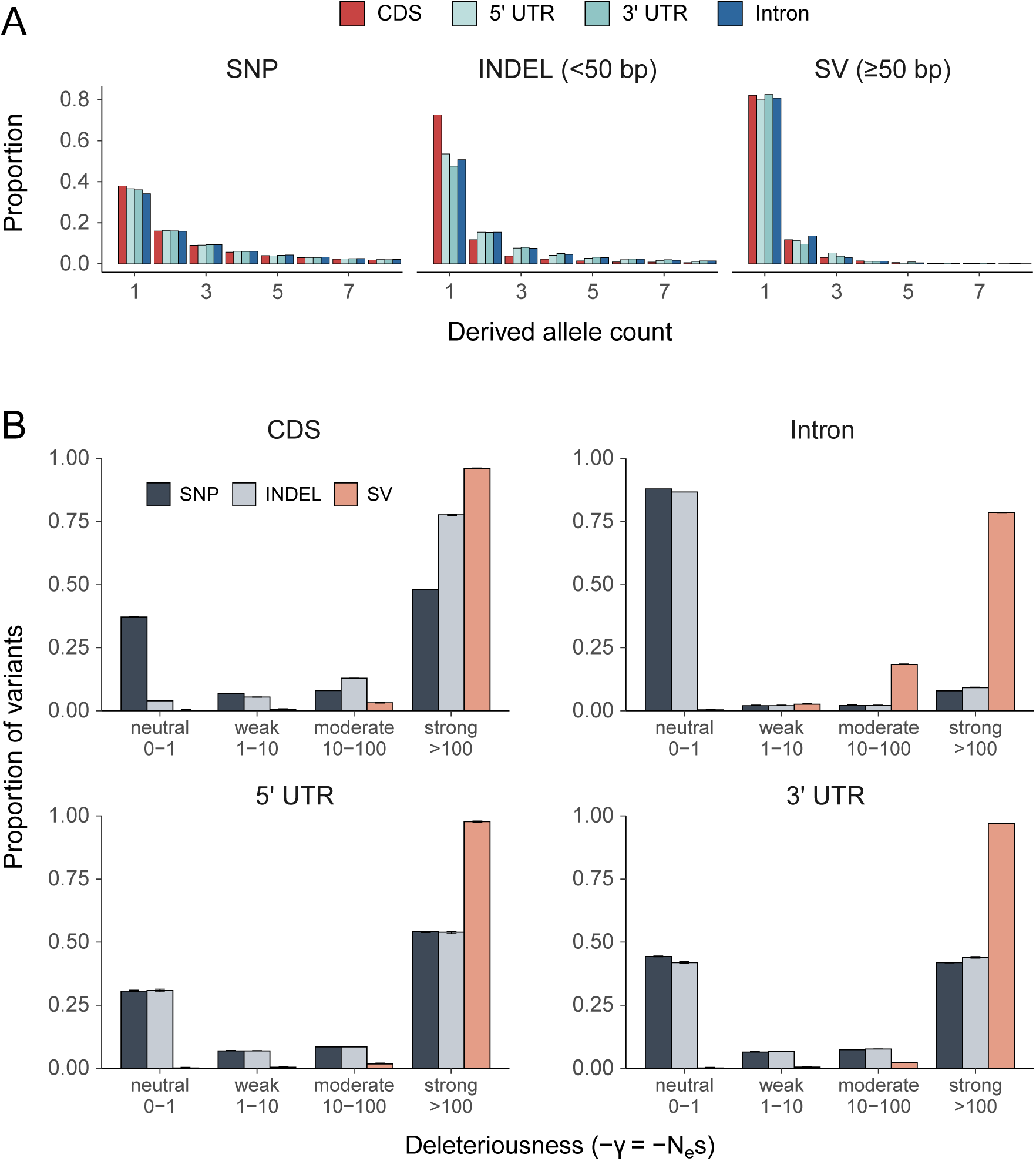
Fitness effects of genome-wide structural variants and single nucleotide polymorphisms. (A) Unfolded site frequency spectrum (SFS) of SNPs, INDELs and SVs residing in genomic regions, CDS, intron, and UTR. The figure displays derived allele counts (DAC) ranging from 1 to 8, with DAC 1 to 31 detailed in SI Appendix, Fig. S6. (B) Distribution of fitness effects (DFE) in bins of population-scaled selection coefficient (γ=*N_e_*s), reflecting variant deleteriousness, a function of the effective population size (*N_e_*) and the selection coefficient (s). DFE was inferred from SFS of different variant types residing in different genomic categories; a similar pattern was yielded by analyses using both fastDFE (as shown) and anavar (SI Appendix, Fig. S7 and Methods). See SI Appendix, Fig. S8 for SFS and DFE per population. Error bars indicate 95% confidence intervals.

The SFSs revealed that all variant types, including SNPs, are predominantly concentrated at low derived allele frequencies, indicating a high incidence of rare variants. SVs and INDELs segregated at much lower average frequencies than SNPs (Fig. 4A; SI Appendix, Fig. S8). In coding regions, which harbor a higher incidence of rare variants compared to other regions (Fig. 4A), most variants are estimated to be deleterious (Fig. 4B), ranging from weakly to strongly deleterious as defined by selective coefficient bins (*γ* = *N_e_s*; Methods), with the majority of SVs (96%) estimated to be strongly deleterious. Similarly, in the regulatory 3’ and 5’ UTRs, we also found an enrichment of strongly deleterious variants, whereas INDELs tend to be more neutral than in coding regions (Fig. 4B). In intronic regions, SNPs and INDELs are estimated to be predominantly neutral, in stark contrast to SVs, which were predominantly identified as strongly deleterious (79%; Fig. 4B). Both fastDFE (Fig. 4B) and anavar analyses yielded similar estimates of the DFE (SI Appendix, Fig. S9). In sum, long variants are in general more deleterious than short variants across the genome, and variants in coding and regulatory regions tend to be more deleterious than those in non-coding regions. Further investigation into population-specific fitness effects revealed a slight increase in strongly deleterious SNPs, INDELs and SVs in the eastern compared to the western population, suggesting that deleterious variants have accumulated detectably in the population with a smaller effective population size (SI Appendix Fig. S10), as predicted by theory (78, 79).

### Confirming and genotyping an 11 megabase pericentric inversion

We were interested in the evolutionary impact and fitness contributions of the largest SV identified, the 11 Mb pericentric inversion. This inversion was confirmed at the HiFi contig level using reference-based alignment (Methods), including dot plots and synteny plots (Fig. 5B-C; SI Appendix Fig. S4), visualization of the inversion subgraph using odgi viz (31) (Fig. 5E), and analysis of gene arrangements across the inversion for haplotypes (Fig. 5F; Methods). The large inversion breakpoints were enriched with repeats, and one breakpoint occurred near the centromere, where segmental duplications and LTRs were also enriched (Fig. 5D). To genotype the inversion in a larger population sample, we performed population genomic analyses on 135 individuals from across the geographic range and across different times since encountering the pathogen at the population level (SI Appendix, Table. S1). The dataset comprised reduced-representation sequencing (RAD seq) data for 108 samples retrieved from Shultz et al. 2016 (74), whole-genome short-read sequencing (WGS) data newly generated for 9 samples (∼15×), and the HiFi data for the 18 pangenome samples (Fig. 6A). The RAD-seq dataset included additional outgroup species, including two Cassin’s Finches (*Haemorhous cassinii*), two Purple Finches (*Haemorhous purpureus*).

**Fig. 5.**
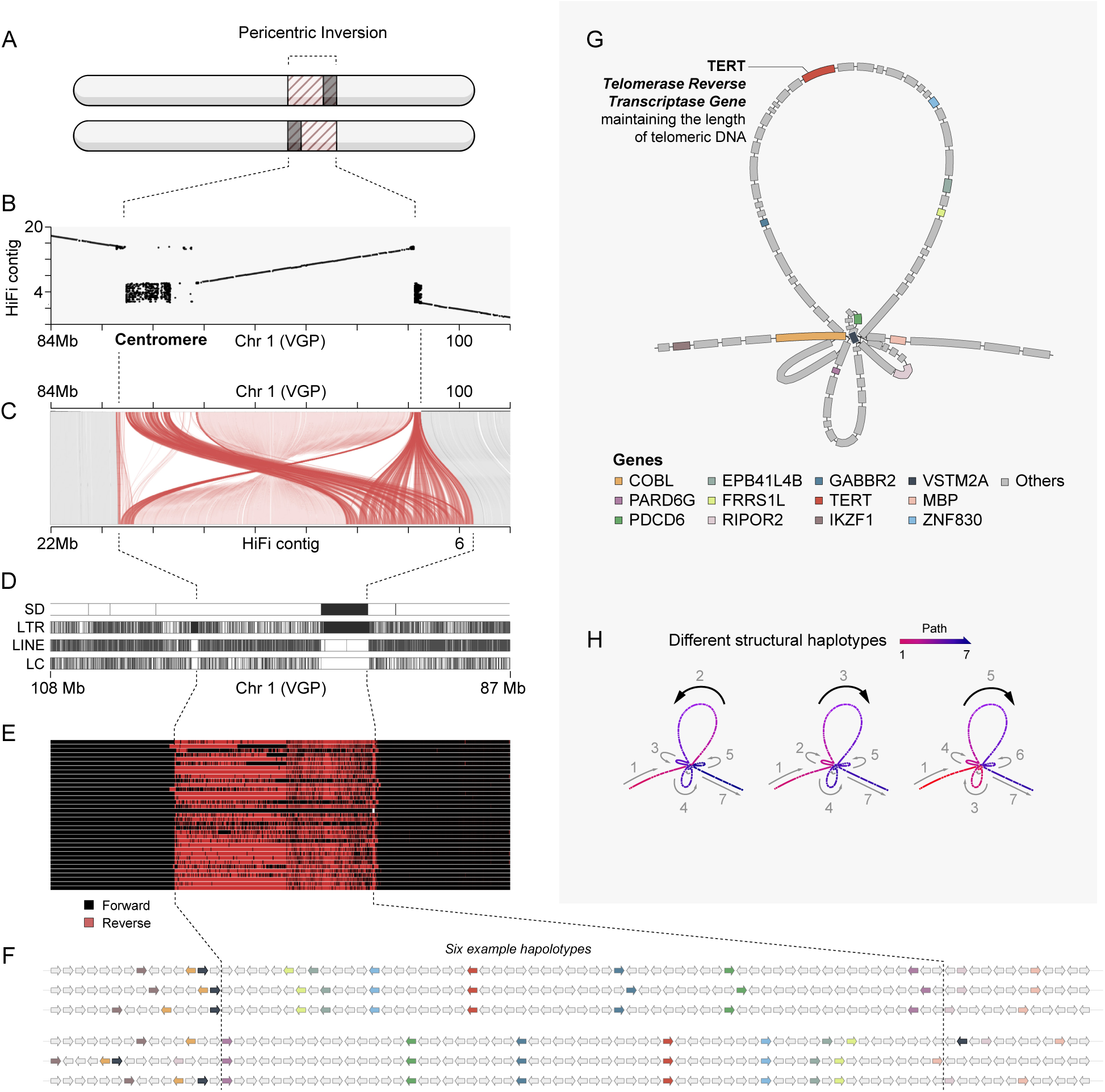
A pericentric inversion on chromosome 1. (A) A schematic of the pericentric inversion. (B) Dot plot of the inversion, displaying the alignment of an NY_2_hap2 haplotype contig (’query’, y-axis) with the VGP genome (’reference’, x-axis). The inversion spans from 89 Mb to 98 Mb on chromosome 1. The presumed centromere, composed of segmental repeats, is subsumed by the inversion. (C) Linear synteny plot of the inversion, correlating with the dot plot (B). (D) Segmental duplications (SDs) and repeat content surrounding the breakpoints. LTR, long terminal repeat; LINE, long interspersed nuclear elements; LC, low complexity. (E) Pangenome graph constructed with PGGB and visualized with odgi viz, depicting flips in strand across the inversion. (F) Genes within the inversion and nearby breakpoints, with those highlighted in color showing differential expression in response to MG infection, were identified through published experimental infection studies and manual searches for immune function (Results). Detailed gene arrangements for all 32 haplotypes are provided in SI Appendix, Fig. S14. The key to these genes is shown in (G), where a pangenome gene graph displays the arrangement of genes within and surrounding the inversion. (H) Examples of haplotype paths among the 32 haplotypes, indicating different structural haplotypes around the inversion and its breakpoints.

**Fig. 6.**
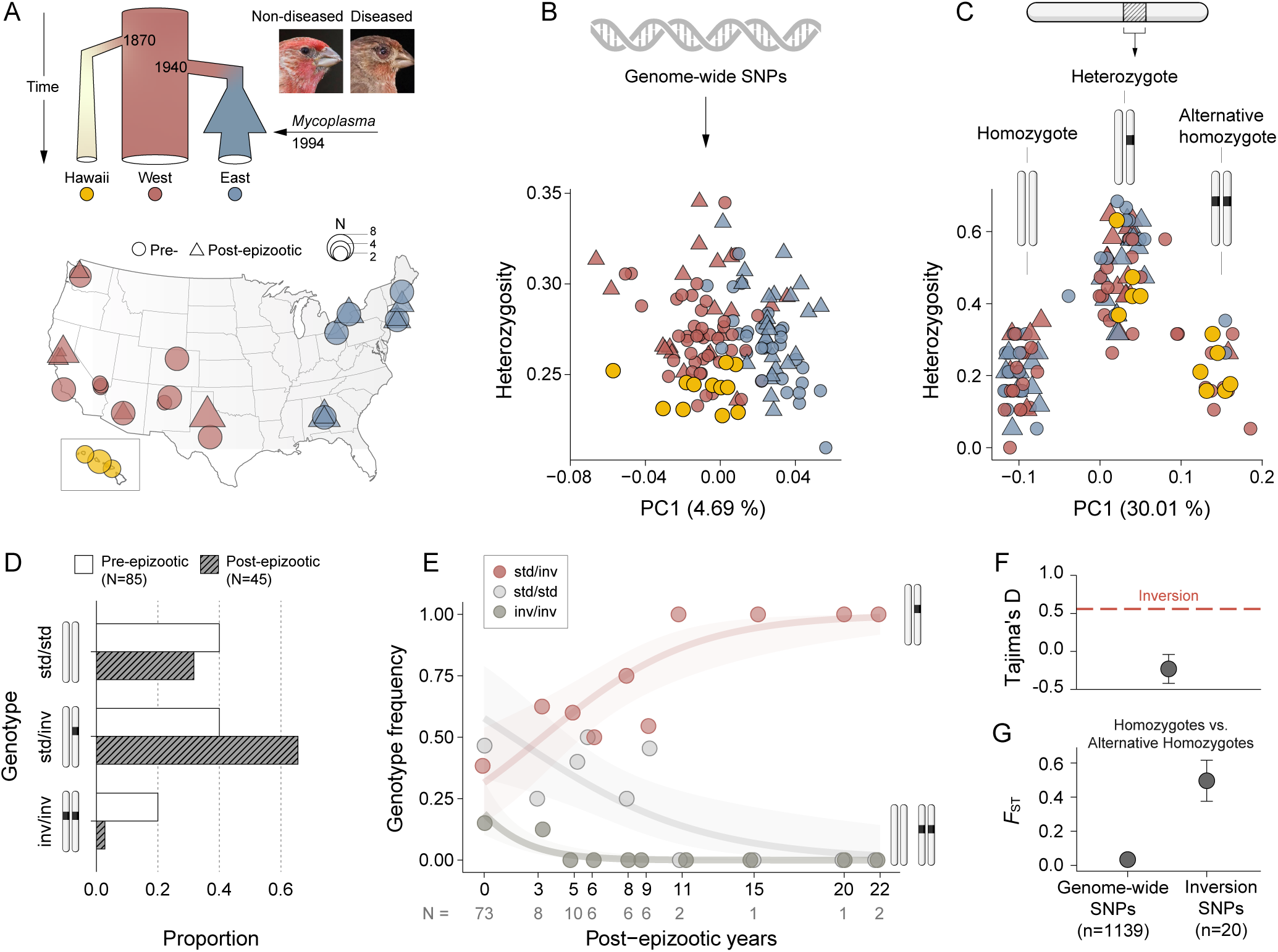
Association between inversion genotype frequency and time since pathogen exposure. (A) Demographic history and map of genetic sampling of House Finch populations, based on data from HiFi sequencing in this study, Shultz et al. (2016; RAD sequencing), and newly sequenced samples (WGS; see Methods). Demographic plot adapted from Shultz et al. (1). (B) PCA and individual heterozygosity based on genome-wide SNPs. The X-axis shows Principal Component 1 (PC1), accounting for the most variance. (C) PCA and individual heterozygosity based on SNPs within the inversion, indicating three inversion genotypes: heterozygote, homozygote, and alternative homozygote, with homozygote as the ancestral type. For inferring the evolutionary history of the inversion, refer to SI Appendix, Fig. S10. (D) Inversion genotype shifts pre- and post-epizootic, suggesting a heterozygote advantage post-epizootic. (E) Genotype frequency of the inversion shifts over the course of the epizootic. Birds from Hawaii, with only pre-epizootic data, are not considered in D and E. (F) Tajima’s D values for the inversion (red line) and chromosomes (mean ± S.E.). (G) Allelic differentiation (*F*_ST_; mean ± S.E.) between individuals with two different homozygous inversion, and between pre- and post-epizootic individuals.

Principal component analysis (PCA) based on 1,159 genome-wide SNPs called from the dataset clustered the 135 individuals by lineage (western, eastern, Hawaii; Fig. 6B; Methods). However, PCA on 20 inversion-associated SNPs formed three distinct clusters with higher heterozygosity in the middle cluster (Fig. 6C). These three clusters thus reflect the three inversion genotypes: homozygote, heterozygote, and homozygote of the alternative arrangement (6). The ancestral haplotype of the inversion was determined to be the haplotype of the homokaryotype identified in the Common Rosefinch (Method; SI Appendix, Fig. S11). The inversion haplotype was found in all three species in the genus *Haemorhous* -- the House Finch, Cassin’s Finch, and Purple Finch -- suggesting pre-existing genetic variation in prior to the origin of House Finches. Thus, the minimum age of the derived inversion is approximately 10 million years (My), which is the time to the most recent common ancestor (TMRCA) of the genus *Haemorhous* (64, 65, 80).

### Heterozygotes for the large inversion have increased in frequency during the *Mycoplasma* epizootic

We found that the inversion genotype frequency shifted significantly between pre- and post-epizootic birds. Specifically, the frequency of the heterozygotes (standard/inversion [std/inv]) increased by 62.5%, from 0.4 to 0.65 (Fig. 6D) across pooled pre- and post-epizootic populations. The frequency of both homozygotes (std/std and inv/inv) decreased, with the inv/inv decreasing by 90% from 0.2 to 0.02 (Fig. 6D). We found that the frequency of the heterozygote (std/inv) genotype increased during the duration of the epizootic across pooled populations (Fig. 6E) and in eastern and western populations separately (SI Appendix, Fig. S12), reaching fixation after 11 years post-epizootic, equivalent to an inversion allele frequency of 0.5. The inversion presents a positive Tajima’s D value (0.56) (81) which surpasses the average genome-wide value (-0.22) (Fig. 6F). Additionally, the inversion exhibited some of the highest population differentiation as measured by *F*_ST_ between individuals with two different homozygotes (Fig. 6G; SI Appendix, Fig. S13). However, we found that population differentiation of the inversion before and after the epizootic is lower than other genomic regions (SI Appendix, Fig. S13), consistent with balancing selection diminishing population differentiation (82). These results suggest that the inversion confers a fitness advantage, because heterozygotes are maintained as a balanced polymorphism within the population, rather than one or the other inversion allele becoming fixed by drift or selection.

We investigated the genic content of the inversion and the diversity of inversion-associated haplotypes in the pangenome graph. Two recent studies, by Veetil et al. (83) and Henschen et al. (54), identified ca. 1780 genes differentially expressed in immune responses of House Finches to MG using experimental infections, suggesting a number of candidates for involvement in the inversion. Of the infection-responsive genes, five (GABBR2, ZNF830, EPB41L4B, FRRS1L, and PDCD6) were found to reside in the inversion, and five (IKZF1, PARD6G, VSTM2A, COBL, and MBP) were found near the inversion breakpoints. Additionally, by searching the NCBI database, we found two immune-related genes (RIPOR2 and RIPK2) near the breakpoints, and a telomerase reverse transcriptase gene (TERT) in the middle of the inversion, which plays a crucial role in the maintenance and lengthening of telomeres (Fig. 5F, G). The functions of these genes and references are listed in the SI Appendix, Table S7. To examine the positional diversity of these genes across haplotypes spanning the inversion, we examine the pangenome gene graph, which confirmed the inversion and identified different haplotype paths through it (Fig. 5F– G). These results indicate dynamic rearrangements of synteny and genes involving genes of immunity and telomerase functions in and surrounding the inversion, potentially contributing to adaptive disease resistance.

### Telomere length decreases with longer times of population exposure to mycoplasmal pathogen

The discovery of the TERT gene within the inversion prompted our interest in variation in telomere length in the context of the epizootic. To this end, telomeric repeats were quantified in the 16 long-read individuals using seqtk telo (84), searching for the (TTAGGG)n repeats in the raw HiFi sequencing reads, thereby circumventing potential assembly-induced biases in telomere reconstruction (Methods). We normalized autosomal read data to a sequencing depth of 20× for consistent comparison across individuals. Our analyses revealed a negative correlation between total telomere length in reads and time elapsed since an individual’s population first encountered MG (Fig. 7), suggesting a potential influence of pathogen exposure or the inversion frequency shifts on telomere dynamics.

**Fig. 7.**
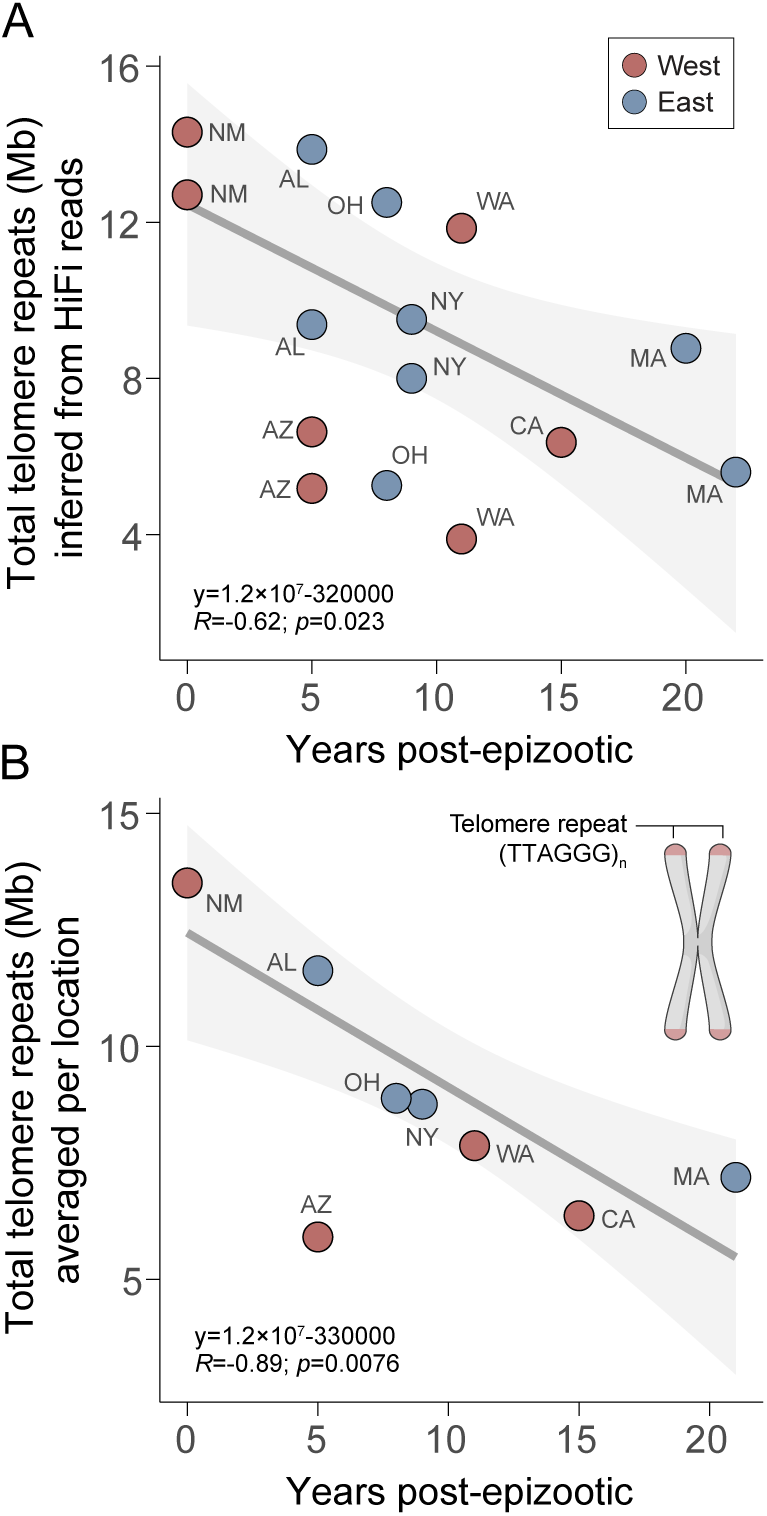
Telomere length decreases over disease exposure history. (A) Scatter plot shows telomere repeat sizes (TTAGGG)n inferred from HiFi reads versus post-epizootic years per sample, with results averaged by location in (B). Regression lines demonstrate a negative correlation between post-epizootic years and telomere repeat sizes, with a 95% confidence interval in shade. HiFi reads are standardized to 20X sequencing depth per sample, and sex-linked reads are excluded. Arizona (AZ) birds displaying severe bill pox, a distinct stressor from the *Mycoplasma* epizootic, are excluded from the regression analysis but are included in the display.

## Discussion

Using long-read sequencing and pangenome approaches on a panel of House Finches, we estimated the fitness consequences of genome-wide variants, including SVs, which have been underrepresented in short-read sequencing data. We detected the imprint of the human-induced introduction of House Finches into the eastern US, including a reduction of genetic diversity for both SNPs and SVs, as well as increased signatures of inbreeding in the introduced eastern population. We also found shorter telomeres in individuals from populations that had been exposed to the *Mycoplasma* pathogen for longer periods of time. Finally, we detectd an association between the largest SV, an 11 Mb pericentric inversion, and time since population exposure to the pathgogen, with a signature of balancing selection in inversion genotype frequencies.

The genome-wide assessment of variants revealed that the vast majority of SVs are estimated to be strongly deleterious across the genome, more so than smaller variants like SNPs and INDELs. This trend is observed across coding, non-coding, and regulatory sequences such as 3’ and 5’ UTRs. Different studies suggest that the fitness effects of SVs arise from both direct and indirect mechanisms. Direct effects include disruption of gene function by altering coding regions, regulatory elements, or by inducing position effects, changing gene dosage, or influencing gene expression at breakpoints (3, 85–87). Indirectly, SVs can affect fitness by suppressing recombination (88, 89). Our findings reveal that the size of SVs relates to their harmfulness, with smaller INDELs being more neutral in non-coding regions compared to larger SVs (Fig. 4B). Given the outlined mechanisms, it is likely that larger SVs have a more pronounced impact on fitness by disrupting gene functions. Empirical studies suggest that a population bottleneck could impact deleterious variation in complex ways, either increasing or reducing the genetic load (79). We found tentative evidence that the eastern population is enriched with more strongly deleterious variants of all types than the western population, suggesting that even a slightly lower effective population size could weaken the efficacy of purifying selection, thus increasing the accumulation of deleterious mutations (90–92). Our findings highlight the necessity of including SVs in studies of genetic diversity and fitness effects to fully understand the evolutionary dynamics of bottlenecks and other demographic events.

Beyond the overall detrimental effects of SVs, our study improves understanding of the potential roles and mechanisms of SV in adaptation. We highlighted a large inversion whose genotype frequencies correlated strongly with population time to exposure to the *Mycoplasma* pathogen, and therefore potentially associated with adaptive evolution in response to disease. This inversion polymorphism is found across all three *Haemorhous* species, with an estimated minimum age of 10 Mya, pointing to the role of standing genetic variation in adapting to environmental challenges, such as pathogen resistance. Theory predicts that recombination suppression and breakpoint mutation (“breakpoint-mutation” theory”) can act together to establish and maintain an inversion (6, 7, 87, 93–97). We found empirical evidence that SVs potentially affect gene expression in genes near breakpoints related to the House Finch immune response (Fig. 5F, G), supporting the “breakpoint-mutation” theory, which suggests that inversions could be selected due to advantageous mutations near breakpoints (93). Overall, our results suggest a versatile role of SVs in influencing the expression of adjacent genes, which could decrease or increase an organism’s fitness depending on gene functions.

Using expression evidence from published studies, we also identified variable structural rearrangements encompassing genes, including the TERT gene, within and nearby the inversion across the 32 haplotype assemblies. Such rearrangements may impact telomerase and immune functions and suggest a putative position effect on genes within and surrounding the inversion due to shifts in their genomic location or surrounding chromatin environment. However, it is important to note that our study did not investigate the mechanisms underlying the balanced polymorphisms of the 11 Mb inversion. Potential factors contributing to this balance could include overdominance, associative overdominance, frequency dependence, and epistasis (5, 7, 8, 95, 97, 98). To gain a deeper understanding of the genetic architecture of the inversion and the mechanisms of its balancing selection, further increasing marker density at the population level, such as through WGS data, and experimental studies, are necessary.

Large inversions have been identified as being associated with avian morphs, behavior, and mating strategies (99). The largest 11.3 Mb inversion we identified is comparable in size to functional inversions in other avian systems, such as a 7.4 Mb inversion in Chickens (*Gallus gallus*) and a 4.5 Mb inversion in Ruffs (*Philomachus pugnax*) (16, 100). Larger inversions have been also observed in other birds, including a 55 Mb inversion in Redpoll Finches (*Acanthis* spp.) (101), a 115 Mb inversion in Quail (*Coturnix coturnix*) (102), and a >100 Mb inversion in White-throated Sparrows (17). However, birds generally exhibit fewer and smaller inversions compared to mammals such as humans and the deer mouse (*Peromyscus maniculatus*), North America’s most abundant mammal. The human genome contains over 1,000 inversions, including >100 large ones (103, 104), while the deer mouse has 21 large inversions and potentially thousands of smaller ones in its genome (10, 105). This suggests that avian inversions are generally shorter and less frequent than in mammals, possibly due to their more streamlined genomes (106), which contain fewer repeats and transposable elements driving the formation of inversions (99, 105).

We observed telomere shortening with prolonged time of population exposure to MG. Extrinsic stressors, including infection and adverse environments, have been reported to drive telomere shortening, potentially involving metabolic, hormonal, and immune mechanisms (107–110). Our findings are consistent with this hypothesis. For example, inside the inversion we identified the TERT gene, encoding the catalytic subunit of telomerase responsible for telomere maintenance and regulating the replicative lifespan of T cells, which is crucial for adaptive immunity (111).

However, the causality between the inversion harboring the TERT gene and telomere shortening remains to be established. Unfortunately, we were unable to include age as a factor in our analyses due to the absence of age information for the birds studied. Telomere length can also be associated with aging (112), although it can also reflect immune function independent of age, as reported in a wild purple-crowned fairy-wrens (*Malurus coronatus*) (113). To minimize the impact of age on observed telomere patterns, we averaged the telomere repeat sizes across birds within each locality. However, future work collecting and incorporating detailed age data into the assessment of telomere shortening is needed.

Certain limitations must be acknowledged to fully interpret our findings. Firstly, the relatively small sample size (16 individuals) cannot fully capture the diversity of the pangenome for all House Finches. Although our evaluation of pangenome graphs suggests a near plateau in the discovery of new core variation and genes using 32 haplotypes, we have potentially overlooked rare genetic variants. Nevertheless, previous studies indicate that more than 8 haplotypes/alleles from a randomly mating population are sufficient to have a high probability of capturing the root of the coalescent tree, which is a major determinant of genetic diversity (114, 115). Secondly, using an outgroup that diverged 12.9 million years ago could affect our ability to accurately deduce ancestral states and derived allele frequencies, which are the basis for assessing fitness effects. To mitigate this, we have employed rigorous criteria for polarizing sites to only homologous outgroup genotypes (Methods) and using anavar, which models and accounts for errors in ancestral state polarization (77). These measures enhance the reliability of our conclusions, despite the challenges of relying on outgroup-based ancestral state reconstructions.

This study underscores the use of combining population-scale long-read sequencing with pangenomic methods in molecular ecology. This approach offers a more complete exploration of genetic variants than is possible with short-read data alone. Other potential approaches to cataloging SVs in natural populations include mapping short reads from many individuals to a pangenome graph comprised of a few individuals (116). This approach could be a favorable economic alternative to sequencing all individuals with long-read methods as we have done here. Looking ahead, the fusion of long-read and pangenomic approaches promises to unveil novel findings about genetic diversity and evolutionary adaptation.

## Materials and Methods

### Sampling and Sequencing

For pangenomic analysis, we retrieved 16 House Finch individuals (SI Appendix, Table S1), with equal representation from the western US (CA, WA, AZ, NM) and eastern US (NY, MA, OH, AL), and a Common Rosefinch individual serving as the outgroup, accessioned in the Museum of Comparative Zoology (MCZ) at Harvard. We isolated high-molecular-weight DNA (HMW DNA) from these samples (muscle or blood) using Qiagen MagAttract HMW DNA Kit and checked the DNA quality using TapeStation (Agilent), Qubit, and NanoDrop (Thermo Fisher Scientific) at the Bauer Core Facility at Harvard. Pacific Biosciences (PacBio) highly accurate long-read (HiFi) sequencing was performed at the University of Delaware Sequencing and Genotyping Center on Sequel IIe SMRTcells. We estimated each population’s exposure time to *Mycoplasma* by calculating the difference between the collection year and the pathogen’s earliest documented arrival from research literature. Detailed sample data is listed in SI Appendix, Table S1. For the 135 individuals used for populations genomic analyses on the inversion (following sections), we obtained double-digest restriction site-associated sequencing (RAD-seq) data, including 104 House Finches with various demographic histories, two Cassin’s finches, and two Purple finches, from Shultz et al. 2016 (74). Additionally, we performed whole-genome resequencing (WGS) at approximately 15x sequencing depth for nine House Finch samples collected from TX and MA, accessioned in MCZ (SI Appendix, Table 1). Their DNA was extracted using Qiagen DNeasy Blood and Tissue kit and sequenced using an Illumina NovaSeq S4 2 x 150 single lane at the Harvard Bauer Core Facility (SI Appendix, Methods).

### Genome Assembly and Annotation

We generated two *de novo* haplotype genome assemblies for each of the pangenome samples using Hifiasm (66). Assembly metrics were assessed with assembly-stats (https://github.com/sanger-pathogens/assembly-stats), and completeness was benchmarked using BUSCO scores (117) (SI Appendix, Table 2). A chromosome-level House Finch genome from California, assembled by the Vertebrate Genomes Project (VGP), was included in our pangenome to provide stable genomic coordinates for downstream analyses. We annotated the VGP genome using approaches based on (i) long-read and short-read RNA-seq data, (ii) homology information, (iii) *ab initio* gene prediction methods, and (iv) gene prediction from projection. We characterized repetitive elements and identified segmental duplications for the VGP genome using RepeatMasker (70) and BISER (69), respectively. To measure telomeric repeats across House Finch samples, we analyzed raw HiFi reads using seqtk telo (84) to avoid assembly biases. Reads linked to sex chromosomes were removed. Each sample was normalized to a 20× sequencing depth with seqtk, and seqtk telo (84) was performed on the normalized autosomal reads. We excluded two low-coverage CA samples (<20× sequencing depth) but supplemented CA data with HiFi reads from the VGP sample, analyzing a total of 15 individuals. We also averaged telomere counts by location by first pooling HiFi reads per location, then performing normalization and analysis with seqtk telo. (SI Appendix, Methods).

### Pangenome Graph Construction and Variant Decomposition

We employed the PGGB pipeline to construct pangenome graphs per chromosome. This approach, validated in projects like the Human Pangenome Project, enables a comprehensive analysis of genetic variation at a resolution that includes SNPs and larger structural variants (INDELs and SVs) (see method selection in SI. Appendix, Methods). We also employed Minigraph (28), a pangenome graph builder by aligning a query sequence against a graph progressively based on the minimap2 algorithm (118), for comparison, and ran Pangene (71), a pangenome gene graph builder, to assess genic variation at gene level. Structural variants and SNPs were decomposed (called) from the graphs using established tools, including vg (119), vcfwave and vcflib (120), bcftools (121) (SI. Appendix, Methods). Following standard definitions, variants were classified based on allele sizes: SNPs if both reference (REF) and alternative (ALT) alleles were 1 bp; multiple nucleotide polymorphisms (MNPs) for allele lengths between 1 and 50 bp; insertions (INS) and deletions (DEL) for size differences under 50 bp; and as SVs (SV-INS and SV-DEL) for differences over 50 bp. Variant polarization was based on the ancestral allele (outgroup). Complex types and multiallelic variants were also categorized (SI Appendix, Methods). autosomal reads. We also averaged telomere counts by location by first pooling HiFi reads per location, then performing normalization and analysis with seqtk telo. (SI Appendix, Methods).

### Identifying and genotyping inversions

Genome-wide identification of inversions, using either assembly- or alignment-based tools, without the aid of manual curation, continues to pose greater challenges compared to the identification of simpler SVs (122). To enhance inversion discovery, we used SVIM-asm (67), an SV caller for haploid or diploid genome-genome alignments, and SyRi (68), a pairwise whole-genome comparison tool. To compile a comprehensive inversion dataset, we merged inversions detected by these methods using bedtools (123), acknowledging the difficulty of accurately identifying all inversions using a single tool (122). To confirm and visualize inversions, dot plots and synteny plots were created based on the pairwise alignments (PAF) of the haplotypes (HiFi contigs) against the VGP genome produced by minimap2 using custom R scripts (examples in SI Appendix Fig. S4). Inversions considered verified had two identifiable breakpoints coexisting in at least one HiFi contig. All six large inversions (>1 Mb) were manually confirmed in HiFi contigs. For the 11.3 Mb inversion described in the study, the PGGB graph visualization (Fig. 6E) was performed using odgi viz (31) based on the sorted subgraph achieved with odgi sort. To genotype the 11.3 Mb inversion across a wider population of 135 House Finches, we mapped sequencing reads (RAD-seq, WGS, and HiFi) to the VGP genome, allowing for variant calling with high accuracy across datasets. Using SNPRelate (124), we performed principal component analysis (PCA) calculated individual heterozygosity on inversion-associated SNPs, revealing the distinct genotypic clusters expected with large inversions (SI Appendix, Methods). The Tajima’s D values for the inversion and for chromosome containing over 100 SNPs were calculated with PopGenome R package (125) (SI Appendix, Methods).

### Distribution of Fitness Effects

The distribution of fitness effects (DFEs) for SNPs, INDELs, and SVs was estimated using the programs fastDFE (76) and anavar (77). These two maximum likelihood approaches use the site frequency spectrum (SFS) to estimate the population-scaled mutation rate (*θ*=4*N_e_µ*; *N_e_* is the effective population size and *µ* is the per site per generation mutation rate) and shape and scale parameters for a gamma distribution of population-scaled selection coefficients (*γ*=4*N_e_s*; *s* is the selection coefficient). Both approaches attempt to control for the confounding effects of demography and polarization errors following the method of Eyre-Walker et al. (2006) (126). We computed the unfolded SFS from the PGGB VCF for focal sets of variants in different categories of variant size (SNPs, INDELs, SVs) and genomic region (CDS, intron, 5’ UTR, and 3’ UTR regions) using custom R scripts, for western, eastern, and both populations combined. We fitted the corresponding SFS in fastDFE and anavar models, using intergenic site frequency spectra as the neutral reference to control for the confounding effects of demography and polarization error. We present the gamma distributions as the proportion of variants falling into four bins of selection coefficients (γ) representing the scaled selection coefficients of variants: neutral (0 ≤ -*N_e_s* ≤ 1), weak (1 < -*N_e_s* ≤ 10), moderate (10 < -*N_e_s* ≤ 100), and strong (-*N_e_s* > 100), and derived 95% confidence intervals through bootstrapping in fastDFE and permutation in anavar (SI Appendix, Methods).

### Analysis of genetic diversity

Runs of homozygosity (ROH) were identified using PLINK (127) based on SNPs across the autosomes, as recorded in the PGGB VCF, with settings of 50 SNPs per ROH (--homozyg-snp 50), a minimum length of 10 kb (--homozyg-kb 10), a gap tolerance of 300 kb (--homozyg-gap 300), and allowing up to two heterozygous SNPs. The individual heterozygosity based on SNPs, insertions, and deletions was calculated only in regions deemed confident by Dipcall (75), a reference-based variant calling pipeline for a pair of haplotype assemblies. The confidence regions were defined as bases covered by an alignment of ≥ 50 kb with a mapQ of ≥5 from each pair and not covered by other alignments of ≥ 10 kb, as implemented in Dipcall (SI Appendix, Methods).

## Acknowledgments

We thank Geoffrey Hill (Auburn University), Allison Shultz, Jonathan Schmitt, Flavia Termignoni Garcia, and Kathrin Näpflin for their contributions to the House Finch collections at the Museum of Comparative Zoology at Harvard University, on which our study is based. We also extend our gratitude to Amberleigh Henschen (University of Memphis) and the VGP for collecting and assembling the VGP House Finch genome. Our thanks also go to Timothy Sackton, Heng Li, Erik Garrison, Andrea Guarracino, and Danielle Khost for their technical advice and feedback on the study, and to Gabriel David and Jacob Höglund for helpful discussion. We are grateful for the computational resources provided by the FASRC Cannon cluster at Harvard University. The work was funded by funds from Harvard University (to S.V.E.), by Harvard Global Institute Fund and Harvard China Fund (to S.V.E. and B.F.), and by the Finnish Cultural Foundation, grant number 00211290 (to B.F.).

## Data availability

The assemblies, raw HiFi, Iso-Seq, and whole-genome sequencing (WGS) data will be made accessible from the NCBI (PRJNA1101522). Analytical scripts are currently available on GitHub at https://github.com/fangbohao. PGGB graphs for each chromosome, along with variant data, can be accessed on Dyrad (DOI: 10.5061/dryad.hhmgqnkqb).

## Author Contributions

S.V.E. conceived the project; and S.V.E. and B.F. designed and developed the research; B.F. performed research including laboratory work, bioinformatic analyses and visualizations; and B.F. and S.V.E. wrote the paper.

