## Supporting Information for "Fitness consequences of structural variation inferred from a House Finch pangenome"

**This PDF file includes:**

Methods

Figures S1 to S15

Tables S1 to S7

**SI Appendix, Methods**

**Sampling and DNA extraction**

We used 16 House Finch samples in the pangenomic analysis, including eight from the western US (western: CA, WA, AZ, NM; two birds in each state) and eight from the eastern US (eastern: NY, MA, OH, AL). These samples were retrieved from the Museum of Comparative Zoology (MCZ) at Harvard (SI Appendix, Table S1) and were all known to have been collected with isolation of high-quality macromolecules in mind. A Common Rosefinch (*Carpodacus erythrinus*; mczbase.mcz.harvard.edu/guid/MCZ:Orn:364823) served as an outgroup. We isolated high-molecular-weight DNA from either blood or muscle tissues of the samples using the Qiagen MagAttract HMW DNA Kit, following the manufacturer's protocols. Before sequencing, we checked the quality of HMW DNA using TapeStation (Agilent), Qubit, and NanoDrop (Thermo Fisher Scientific) at the Bauer Core Facility at Harvard. We also used a chromosome-level House Finch genome from CA, collected by Amberleigh Henschen (University of Memphis) and assembled by the Vertebrate Genomes Project (VGP; www.genomeark.org/genomeark-all/Haemorhous_mexicanus.html; primary assembly; hereafter referred to as the VGP genome) in this pangenomic study.

For the 135 individuals used for populations genomic analyses on the inversion (following sections), we obtained double-digest restriction site-associated sequencing (RAD-seq) data, including 104 House Finches with various demographic histories, two Cassin's finches, and two Purple finches, from Shultz et al. 2016 (1). Additionally, we performed whole-genome resequencing (WGS) at approximately 15x sequencing depth for nine House Finch samples (blood or muscle tissues) collected from TX and MA, accessioned in MCZ (SI Appendix, Table 1). For this, DNA was extracted from the tissue samples using a DNeasy Blood and Tissue kit (Qiagen, Valencia, California, USA) following the manufacturer's protocols. The transposase-mediated fragmentation-based library protocol was applied, and the DNA was sequenced using an Illumina NovaSeq S4 2 x 150 single lane at the Harvard Bauer Core Facility.

We estimated the exposure time (years) of each population to *Mycoplasma* by subtracting the pathogen's putative arrival year from the collection year. We determined the pathogen's arrival year from the earliest documented research literature in the area or state (if no reports were available in the specific localities we sampled). The SI Appendix, Table S1, lists detailed information about all samples used, including sex, locality (GPS), collection years, putative exposure years, references for estimating exposure years per population, and sequencing depth.

**Sequencing, genome assembly, and genome annotation**

Pacific Biosciences (PacBio) highly accurate long-read (HiFi) sequencing was performed at the

University of Delaware Sequencing and Genotyping Center. Briefly, libraries were prepared using the SMRTbell Express Template Prep Kit 2.0 (PacBio) and the Blue Pippin DNA Size Selection System (Sage Science), and then sequenced on Sequel IIe SMRTcells (PacBio). We converted PacBio BAM files to FASTQ files using bam2fastx (PacBio) and assembled the HiFi reads into two pseudo-haplotype assemblies and a primary assembly at the contig level for each sample using Hifiasm (2). Assembly quality metrics were quantified with assembly-stats (https://github.com/sanger-pathogens/assembly-stats), and assembly completeness was benchmarked by Universal Single-Copy Orthologs (BUSCO) assessment (3). The quality of each de novo genome is reported in the SI Appendix, Table 2.

**Gene annotation**

When this study was initiated, the current NCBI House Finch annotation (GCF_027477595.1-RS_2023_09; 98.9% complete BUSCOs) had not been released. We annotated coding genes for the VGP genome using approaches based on (i) long-read and short-read RNA-seq data, (ii) homology information, (iii) *ab initio* gene prediction methods, and (iv) gene prediction from projection. Specifically, we first performed PacBio long-read RNA sequencing (Iso-Seq) for four tissues (heart, brain, liver, and eye) of a sample from the MCZ (https://mczbase.mcz.harvard.edu/guid/MCZ:Orn:365346) at the University of Delaware Sequencing and Genotyping Center. Transcripts per tissue were assembled from long-read RNA alignments (BAM) using IsoQuant (4), a *de novo* transcript discovery tool for Iso-Seq data. Additionally, we reconstructed a spleen transcriptome from short-read RNA sequences (raw RNA sequences retrieved from Zhang et al. (5)) using StringTie (6). Coding regions (open reading frames) of the assembled transcripts were predicted by TransDecoder (https://github.com/TransDecoder/). The BRAKER2 pipeline (7) was used to annotate coding genes by integrating protein evidence prediction and de novo gene prediction with GeneMark-EP+ (8) and AUGUSTUS (9). For this, available protein sequences of chicken and House Finch were retrieved from NCBI in FASTA format and provided to BRAKER2.

To project gene models from well-annotated genomes to the House Finch genome, we used TOGA (Tool to infer Orthologs from Genome Alignments) (10) and miniprot (11). Using TOGA, we projected annotations of coding genes from Chicken (*Gallus gallus*, GCA_016699485.1), Zebra Finch (*Taeniopygia guttata*, GCA_003957565.4), and Common Canary (*Serinus canaria* GCA_022539315.2). For this, we input pairwise genome alignment chains between the three designated reference genomes and the VGP genome using LASTZ (12). Using miniprot, we aligned protein sequences of the designated reference genomes to the VGP genome for gene annotation. Gene models from all the above prediction approaches were then integrated using StringTie-merge, and duplicates and incomplete structures were filtered out using AGAT (https://github.com/NBISweden/AGAT). We eventually annotated 22,080 protein-coding genes in the VGP genome with 98.6% complete BUSCOs ("protein" mode), a similar quality metric to the NCBI annotation released later in September 2023. We obtained very similar results for analyses based on our and NCBI's annotations, and here report results based on our genome annotation.

**Repeat Annotation**

Repeat annotation on the VGP genome was performed using the RepeatModeler and RepeatMasker pipelines (13). We predicted repetitive elements *de novo* using RepeatModeler through three complementary programs: RECON (14), RepeatScout (15), and Tandem Repeats Finder (16). Satellite Repeat Finder (17) was used to identify satellite DNA regions. We then manually curated repeat libraries by concatenating the Avian Repbase transposable element library (18) and the de novo repeat libraries and ran RepeatMasker with this curated library. To assess the repeat content of SVs, we implemented repeat annotation on the longest allele of each SV site identified (see Methods below), using RepeatMasker based on the same repeat library curated. To identify segmental duplications, we used the masked version of VGP genome derived from RepeatMasker as input for BISER (19) with default parameters.

To measure telomeric repeats in each individual, we used seqtk telo (20) directly on raw HiFi reads to avoid assembly biases. The two CA samples were excluded from this analysis due to their low sequencing depth (~15×), but the HiFi reads of the VGP sample (collected from CA) were retrieved and used to supplement the CA data, resulting in a total of 15 individuals for this analysis. To exclude HiFi reads linked to sex chromosomes, the reads were mapped to the VGP genome using minimap2 (21), and the sex-linked reads were excluded. To normalize the data, HiFi reads per sample were subsampled to a sequencing depth of 20× using seqtk. Seqtk telo (20) was then performed on the normalized autosomal reads. We also averaged telomere counts by location by first pooling HiFi reads per location, then performing normalization and analysis with seqtk telo.

**Pangenome graph construction**

**PGGB.** PGGB is a pipeline for constructing unbiased pangenome graphs using all-to-all genome alignments without relying on a reference genome (22). The PGGB pipeline was selected as the primary pangenome graph builder based on the study's aims and the corresponding merits of the tool. Specifically, i) the PGGB graph represents variants of all sizes, including SNPs, INDELs (<50 bp), and SVs (>50 bp), and stores all coordinates from the input assemblies, providing a comprehensive picture of genomic variation (22-25). This feature aids our downstream comparative population genomic analyses, which include assessing the fitness effects of variants of various sizes; ii) PGGB has been applied and validated on a large scale in pioneering projects such as the Human Pangenome Project (23, 24, 26, 27); iii) PGGB identifies comparable SVs in accuracy and quantity, compared with other mainstream graph builders like minigraph and minigraph-cactus (24, 28-30); iv) This study was less concerned about the computational time and resources for PGGB graph construction, which might be a demerit compared to fast graph builders like minigraph (24, 28, 31). However, the latter does not detect small variants.

We applied PGGB to build a pangenome graph from 32 House Finch haplotype assemblies, the VGP primary assembly, and two Common Rosefinch (outgroup) haplotype assemblies following the pipelines of creating the human pangenome by the Human Pangenome Reference Consortium (23). Prior to running PGGB, we renamed HiFi contigs according to the Pangenome Sequence Naming (github.com/pangenome/PanSN-spec) with fastix (github.com/ekg/fastix), and partitioned them into 39 autosomes (communities) by mapping each assembly against the VGP assembly with wfmash (-s 5000, segment length; -p 90, percent identity). The unmapped contigs were remapped in split mode, and the best mapping for each split rescue per chromosome was retrieved. We applied PGGB to each chromosome separately, in three main steps. First, sequence base-level alignment is performed with wfmash (30) based on the input assemblies. We used default settings including mapping percent identity (-p 90) and segment length (-s 5000). Next, seqwish (30) was used to build pangenome graphs of each community. We set the minimum match length to 79 (-k) during graph induction as advised by PGGB developers. Finally, the graph was refined by smoothxg and gfaffix (32) through compressing and organizing. These parameters are reasonable when applied across the whole genome, but can be refined when applied to individual regions depending on the local sequence divergence, repeat content, and other factors. The pangenome growth curve analysis (Fig. 2A) was performed with panacus (https://github.com/marschall-lab/panacus).

**Minigraph**. Minigraph generates a pangenome graph by aligning a query sequence against a graph progressively based on the minimap2 algorithm (28). Using the VGP genome as the reference assembly, minigraph converted its FASTA format to GFA, serving as the initial graph. Minigraph progressively mapped additional assemblies to the graph and augmented SVs of at least 50 bp in size into the graph incrementally. Minigraph shows some sensitivity to the input order of genomes, especially with the first input sequence acting as a reference or backbone (28, 30). To run Minigraph with the least bias, we used the default -cxggs options and input the assemblies in the following order: VGP genome, 32 House Finch assemblies in a random order, and two Rosfinch assemblies last.

Chromosome 16 of the VGP genome, with 93% of its composition being repeat content, exhibits an unusually high proportion of repeats. This was confirmed by its repeat annotation (SI

Appendix, Table S4), by mapping 32 haplotype assemblies to the VGP genome (see SI Appendix, Fig. S15), and by examining the resulting graph (SI Appendix, Table S4). The PGGB produced exceptionally large graph for the chromosome 16, with large bubbles nesting subgraphs (termed snarls) containing multiple variants that cannot be further decomposed. We refrain from further biological interpretation of chromosome 16 in this study; however, we have deposited its graph for future exploration (Data availability). To reduce false positive variants, such as singletons, chromosome 16 was excluded from the pangenomic and site frequency spectrum analyses.

**Pangene.** Pangene summarized the 32 pairwise miniprot alignments to the VGP reference and produced a pangenome gene graph in GFA format. We aligned proteins from the VGP genome to 32 haplotype assemblies individually using miniprot with the parameter "--outs=0.97" and built the graph using default settings in Pangene. We then decomposed the gene presence-absence and copy number variation from the graph using "pangene.js gfa2matrix". The visualization of the Pangene graph was performed using gfa-server and Bandage (33). $F_{ST}$ of PAV polymorphism was calculated using R package SNPRelate (34).

**Calling genomic variants from graphs**

The PGGB graph, in GFA format, was decomposed into a VCF for each chromosome using vg deconstruct (35). Following protocols of the Human Pangenome Consortium, the large (>100 Kb) SVs in the VCF were removed to avoid spurious variants using vcfbub from vcflib (36) with the options -l 0 -a 100,000. The VCF files were then normalized to decompose complex alleles into primitive ones using vcfwave with the option -l 1,000, and using bcftools (37) with the option “norm -m+both” to join biallelic sites into multiallelic records separately for SNPs and indels. For the minigraph graph (GFA), SVs (> 50 bp) were called using "minigraph -cxasm --call" for each sample in BED files outputted from minigraph. We then merged the per-sample calls, generating a VCF of all chromosomes with “mgutils.js merge”.

**Filtering and classifying SVs.**

We focused on genomic variants within House Finch haplotypes, retaining those called in at least 90% of the 32 haplotypes (29 haplotypes). We polarized the genotype of ancestral and derived alleles in these haplotypes via custom R-scripts, using the outgroup genotype (Common Rosefinch; haplotype 1) as a reference. Polarization was limited to sites where alleles in House Finches matched alleles in homologous Common Rosefinch genotypes (0/0, 1/1, 1/., or 0/..). Variant types were categorized based on the size of reference (REF) and alternative (ALT) alleles. Variants were categorized as SNPs when both REF and ALT lengths were 1 bp (length (REF) = length (ALT) = 1 bp), and as multiple nucleotide polymorphisms (MNPs) when their lengths ranged between 1 and 50 bp (length (REF) = length (ALT) = 1 < 50 bp). Variants were designated as insertions (INS) and deletions (DEL; altogether INDELs) when there was a size difference between the longest and shortest alleles of less than 50 bp, and as SVs when this difference exceeded 50 bp. INDELs and SVs were further classified according to the ancestral allele (ANC). Specifically, deletions (DEL and SV-DEL) were identified at sites where the derived allele was 1 bp in size (length (DER) = 1), and insertions (INS and SV-INS) at sites where the ancestral allele was 1 bp (length [ANC] = 1). Variants that did not fit these criteria were labeled as complex types (INDEL-complex and SV-complex). Additionally, variants with more than two alleles were classified as multiallelic.

**Identifying inversions.**

Genome-wide inversion identification using either assembly- or alignment-based tools, without the aid of manual curation, continues to pose greater challenges compared to the identification of simpler SVs (38). To enhance inversion discovery, we additionally performed SVIM-asm (39), an SV caller for haploid or diploid genome-genome alignments, and SyRi (40), a pairwise whole-genome comparison tool. We called SVs with SVIM-asm for the 32 pairwise alignments (BAM files) produced by minimap2, in diploid mode (svim-asm diploid). The individual SVs in VCF format were merged using Jasmine (41), with the parameters "--max_dist_linear=0.1 --min_dist=50," requiring variant breakpoints to be within a 50 bp window to exclude redundant SVs. SyRi identifies structural rearrangements between genomes using whole-genome alignment based on chromosome-level assemblies (40). Thus, as recommended by SyRi pipeline, we generated 32 pseudo-chromosome-level assemblies using RagTag (42), a scaffolding tool that maps query scaffolds to a high-quality reference genome, here the VGP genome. To compile a comprehensive inversion dataset, we merged inversions detected by these methods, acknowledging the difficulty of accurately identifying all inversions using a single tool (38). This merging was done with bedtools, allowing a maximum distance of 100 bp between inversions for merging.

To confirm and visualize inversions, dot plots and synteny plots were created based on the pairwise alignments (PAF) of the haplotypes (HiFi contigs) against the VGP genome produced by minimap2 using custom R scripts. Inversion breakpoints were identifiable when a HiFi contig covers the inversion (covering two breakpoints) and aligns in reverse within the inversion area compared to the reference genome, or when a contig covers just one breakpoint and maps to the reference in two opposite orientations (examples in SI Appendix Fig. S4). Inversions considered verified had two identifiable breakpoints coexisting in at least one HiFi contig. All six large inversions (>1 Mb) were manually confirmed using HiFi contigs through the methods described. We visualized the 11.3 Mb inversion described in the study from the PGGB graph (GFA) directly using odgi viz (43), based on the sorted subgraph achieved with odgi sort. And we also visualized the Pangene graph to examine the genes and haplotype paths within and surrounding the inversion.

**Genotyping the 11 Mb inversion**

Large-sized inversions can be genotyped with sparse sequencing data, such as reduced-representation sequencing data (44). To genotype the 11 Mb inversion on chromosome 1 across populations with various epizootic histories, we conducted PCA and heterozygosity analyses of the inversion-associated SNPs using the 135 individuals as introduced in "*Sampling and DNA extraction*".

To call consensus SNPs from all HiFi, RAD, and WGS reads, we adopted reference-based mapping approaches for each dataset individually, and then merged and filtered VCF files to obtain a final SNP dataset. The VGP genome, indexed with samtools faidx, served as the reference genome to provide uniform coordinates. For RAD and WGS data, reads were mapped to the reference using bowtie2 (45) with the options --very-sensitive-local -I 149, and the mate-pair information was verified and corrected using Picard tools (https://broadinstitute.github.io/picard). SNPs were called with bcftools mpileup per chromosome with the options -d 10000 -q 30 -Q 30. For HiFi data, raw reads were mapped to the VGP genome using minimap2 with the -ax map-hifi option, and variants were called from the resulting BAM files per chromosome for each sample using Longshot (46), a long-read-specific variant caller. Then, we concatenated chromosomes and merged 135 samples from the above VCF files using bcftools concat and merge.

We filtered the VCF by retaining biallelic SNPs with a minimum of two for minor allele count (-mac 2) and no more than 5% missing data (--max-missing 0.95) using vcftools (47). This stringent filtering ensured well-represented consensus variants across three datasets, resulting in 1,159 SNPs and 20 inversion-residing SNPs across the genome. To genotype the inversion, we used the R package SNPRelate (34) for PCA of inversion-associated SNPs and computed heterozygosity as the percentage of heterozygous sites in the 135 samples in R. PCA of an inversion typically displays three distinct clusters: homokaryotypes of one arrangement, heterokaryotypes, and homokaryotypes of the alternative arrangement, with the middle cluster (heterokaryotypes) showing high heterozygosity. For comparison, PCA and heterozygosity analyses were also conducted on the genome-wide 1,159 SNPs, as shown in Fig. 6B.

**Determining derived inversion haplotype.** We inferred the ancestral (standard) haplotype and the minimum age of the derived haplotype based on the inversion genotype in outgroup species, namely Cassin’s Finch, Purple Finch, and Common Rosefinch, and their phylogenetic relationship with the House Finch (SI Appendix, Fig. S10). We assigned the Common Rosefinch's inversion genotype as the ancestral (standard) arrangement and identified the minimum age of the derived inversion as the time of the most recent common ancestor (TMRCA) of the House Finch, Cassin’s Finch, and Purple Finch, using available phylogenetic history (48-50) (SI Appendix, Fig. S10).

**Distribution of fitness effects**

The distribution of fitness effects (DFEs) for SNPs, INDELs, and SVs was estimated using the programs fastDFE (51) and anavar (52). These two maximum likelihood approaches use the site frequency spectrum (SFS) to estimate the population-scaled mutation rate ($\theta=4N_e\mu$; $N_e$ is the effective population size and $\mu$ is the per site per generation mutation rate) and shape and scale parameters for a gamma distribution of population-scaled selection coefficients ($\gamma=4N_es$; $s$ is the selection coefficient). Both approaches attempt to control for the confounding effects of demography and polarization errors following the method of Eyre-Walker et al. (2006) (53).

We computed the unfolded SFS from the PGGB VCF for focal sets of variants in different categories of variant size (SNPs, INDELs, SVs) and genomic region (CDS, intron, 5’ UTR, and 3’ UTR regions) using custom R scripts. We computed the SFS for western, eastern, and both populations combined. To estimate the DFEs in fastDFE, we fitted the corresponding SFS to the ‘GammaExpParametrization’ model. To estimate the DFE using anavar, we fitted the corresponding SFS to the ‘neutralSNP_vs_selectiveSNP’ model for SNP datasets and the ‘neutralINDEL_vs_selectedINDEL’ model for INDEL and SV datasets, using a continuous gamma distribution. Both programs applied the SFS from a putatively neutral set of variants to control for the confounding effects of demography and polarization error. We used the SFS of variants from intergenic regions (1Kb apart from genes) as neutral references. We present the gamma distributions as the proportion of variants falling into four bins of selection coefficients (γ) representing the scaled selection coefficients of variants: neutral ($0 \leq -N_es \leq 1$), weak ($1 < -N_es \leq 10$), moderate ($10 < -N_es \leq 100$), and strong ($-N_es > 100$). All estimates of $\gamma$ are negative. We obtained 95% confidence intervals for DFE analyses by parametric bootstrapping in fastDFE (inf.bootstrap function) and permutating genes in anavar analyses.

**Calculating runs of homozygosity and heterozygosity**

To investigate the genetic diversity of the western and eastern populations, runs of homozygosity (ROH) were identified based on SNPs across the autosomes, as recorded in the PGGB VCF. ROHs were analyzed using PLINK (54), with settings of 50 SNPs per ROH (--homozyg-snp 50), a minimum length of 10 kb (--homozyg-kb 10), a gap tolerance of 300 kb (--homozyg-gap 300), and allowing up to two heterozygous SNPs. We also calculated individual heterozygosity based on

SNPs, insertions, and deletions in their respective populations. For a strict comparison, heterozygosity was calculated only in regions deemed confident by Dipcall (55), a reference-based variant calling pipeline for a pair of haplotype assemblies. The confidence regions were defined as bases covered by an alignment of ≥ 50 kb with a mapQ of ≥5 from each pair and not covered by other alignments of ≥ 10 kb, as implemented in Dipcall.

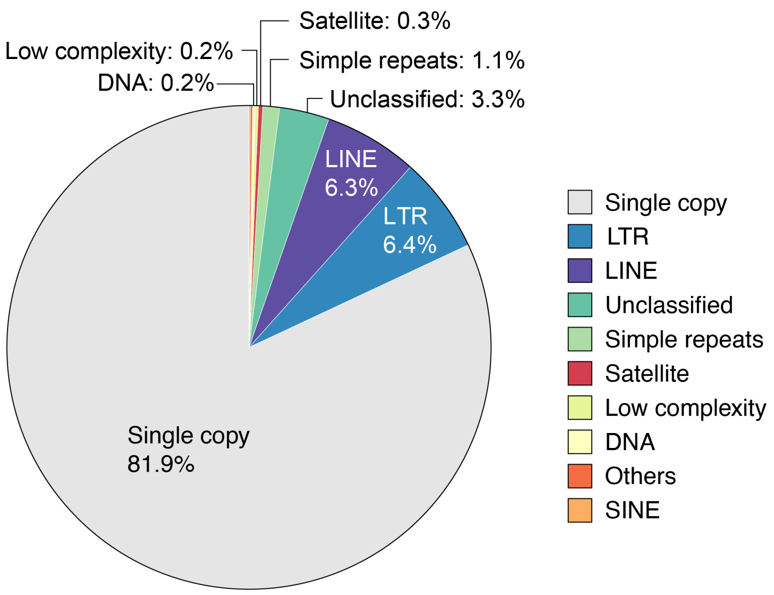


**Fig. S1.** Repeat landscape of the House Finch genome curated from the Vertebrate Genomes Project (VGP genome). See SI Appendix, Table S4 for a detailed report on repeat content of the genome and per chromosome.

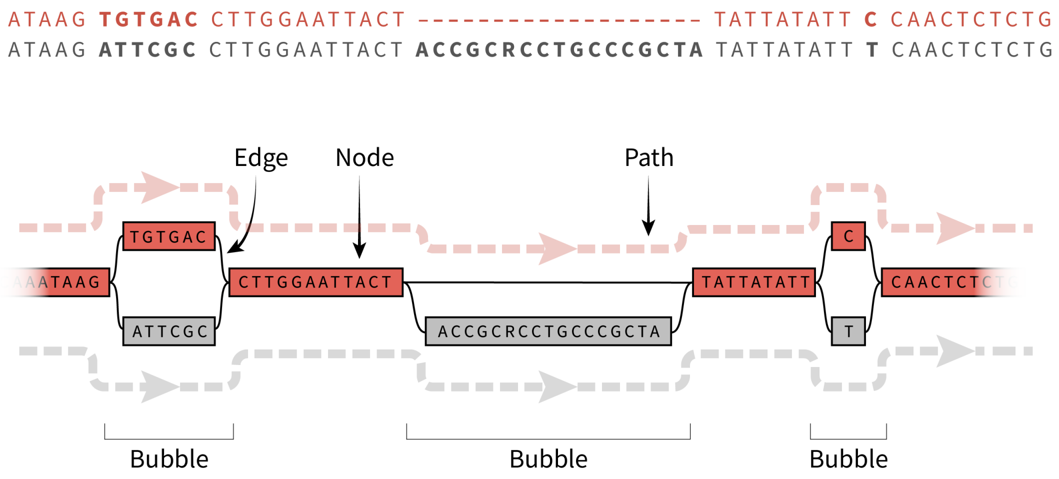


**Fig. S2.** A schematic pangenome variation graph. The graph model comprises a sequence graph, where nodes symbolize oriented DNA segments, and bidirected edges indicate connectivity relationships between these nodes. Nodes can orient in forward or reverse directions, creating a bidirected graph with four possible edges between any node pair to represent all combinations of orientations. Embedded within this graph are haplotype paths (colored dotted lines) representing individual haplotypes. The graph also features 'bubble' structure to characterize variants.

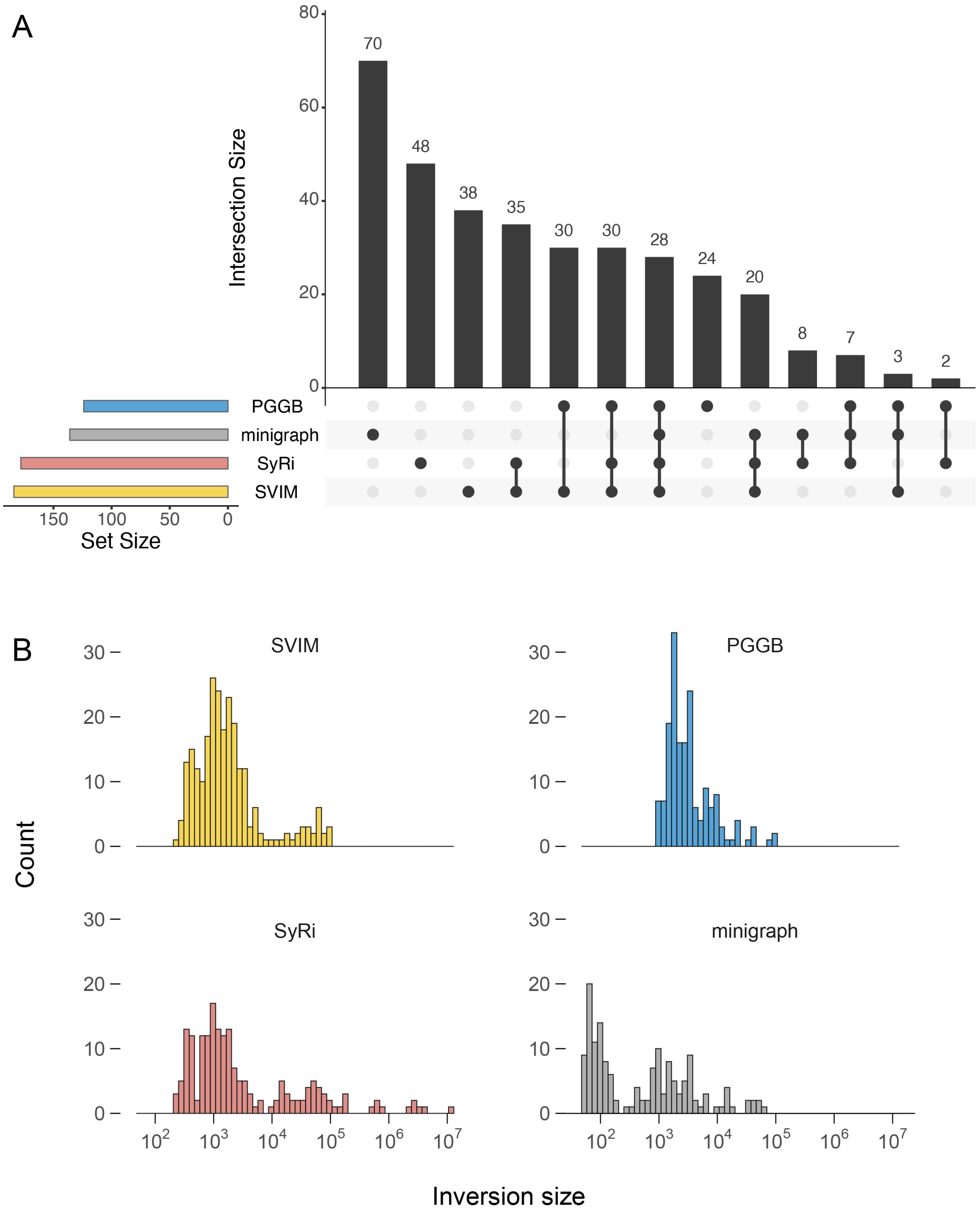


**Fig. S3.** Overview of 343 inversions detected using four different programs. (A) An UpSet plot illustrates the intersection of inversions identified across the four programs. (B) Size distribution of inversions detected by each program. All six large inversions (>1 Mb; identified by SyRi as shown above) were verified using dot plots in HiFi contigs (Methods).

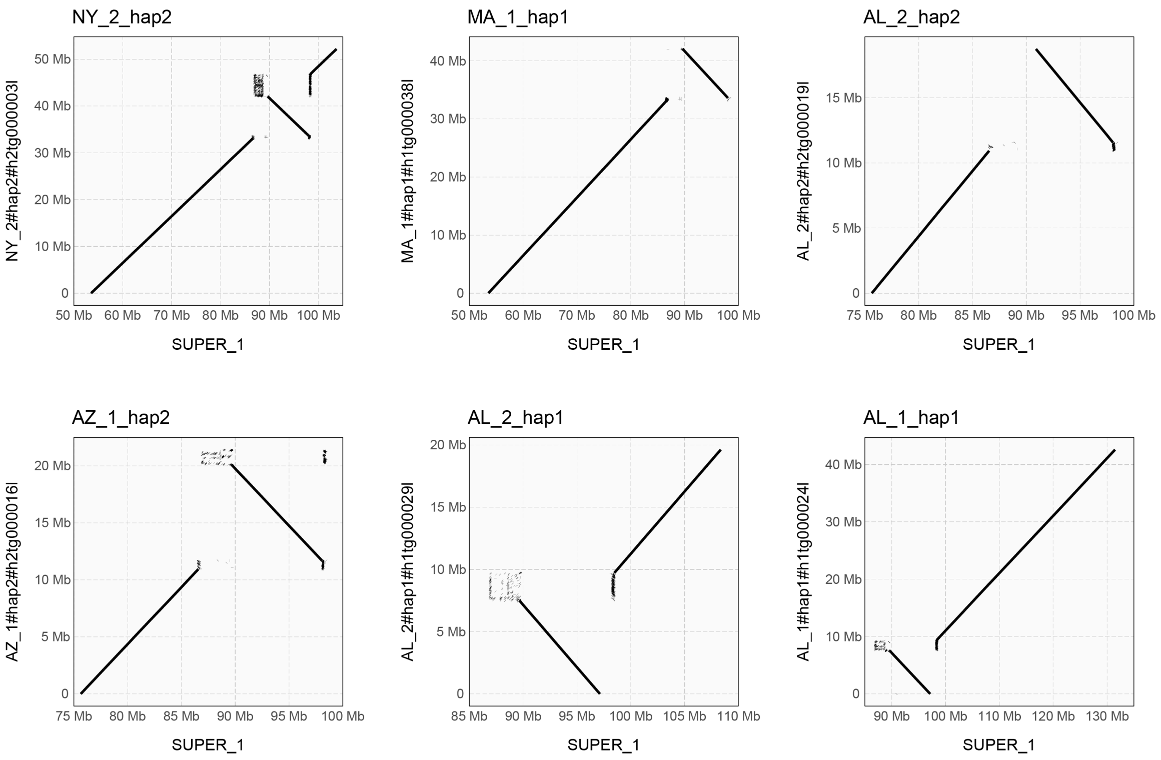


**Fig. S4.** Dot plots illustrating the 11.3 Mb inversion on chromosome 1, extending from 87 Mb to 98 Mb. The Y-axis represents the haplotype HiFi contig ('query'), and the X-axis corresponds to the VGP genome ('reference').

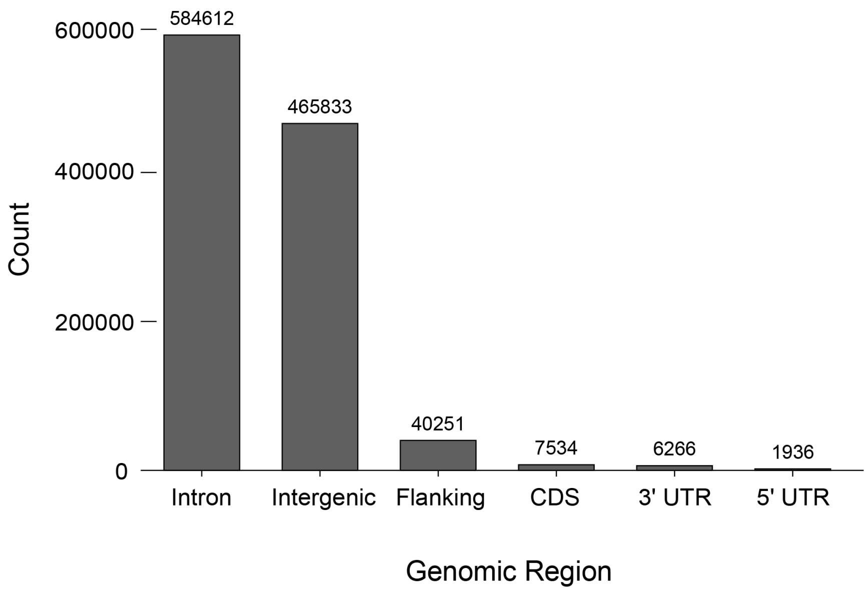


**Fig. S5.** Frequency of structural variants (SVs) in various genomic regions.

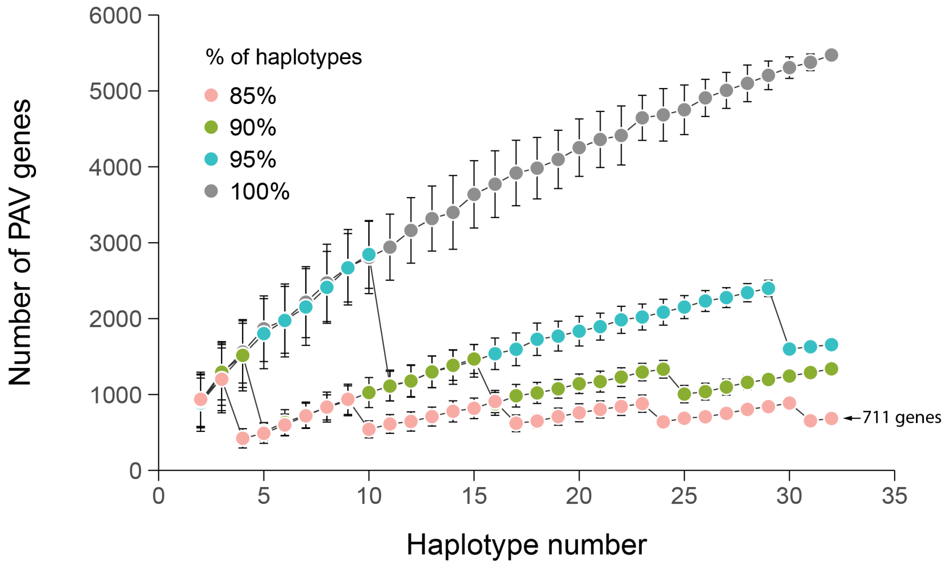


**Fig. S6.** Gene presence-absence variation derived from the pangenome gene graph constructed with miniprot and pangene (Methods). To reduce false positive gene variation caused by fragmented HiFi contigs, which Pangene filters out and thus flags as potential gene absence, and due to the challenges of mapping protein sequences to genomes (11, 56).

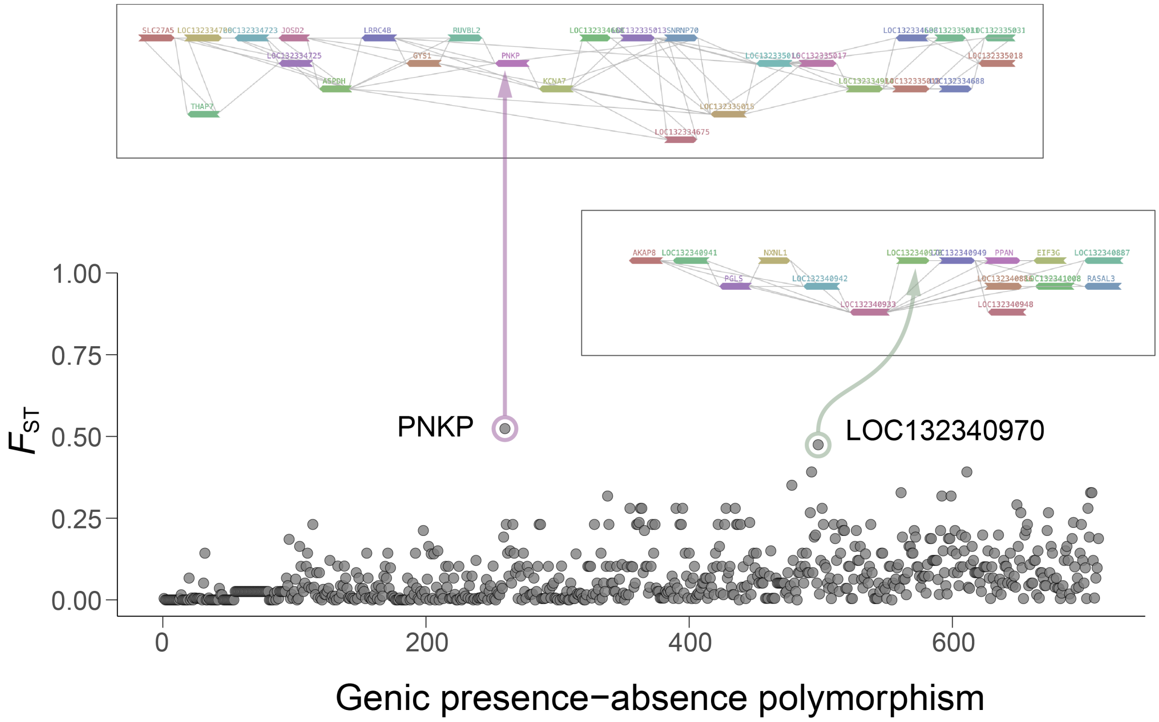


**Fig. S7.** Allelic differentiation ($F_{ST}$) of PAV genic polymorphisms. Outliers ($F_{ST}$ > 0.4) were manually inspected using Pangene visualizations (Methods). as potential false positives due to fragmented contigs.

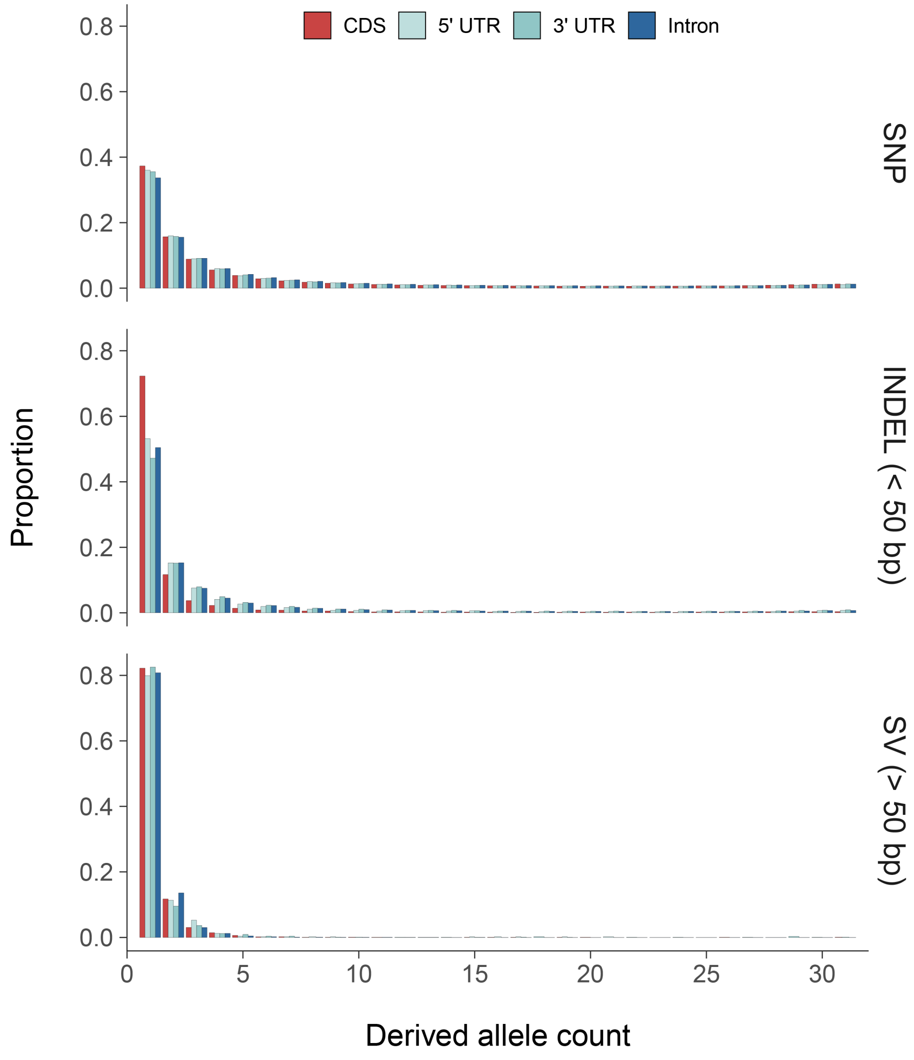


**Fig. S8.** Unfolded site frequency spectrum (SFS) of SNPs, INDELs and SVs residing in genomic regions, CDS, intron, and UTR.

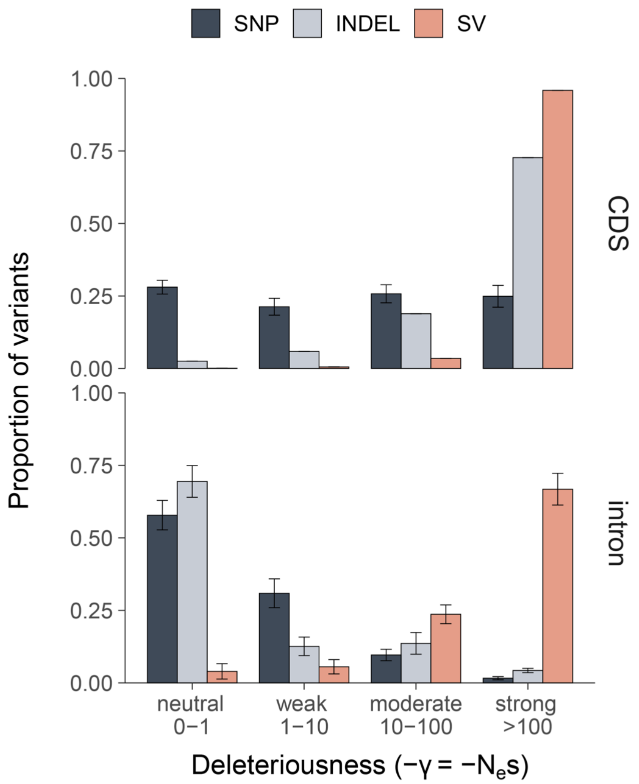


**Fig. S9.** Distribution of fitness effect (DFE) resulted from Anavar program, in bins of population-scaled selection coefficient (γ=*Ne*s), reflecting variant deleteriousness, a function of the effective population size (*Ne*) and the selection coefficient (s).

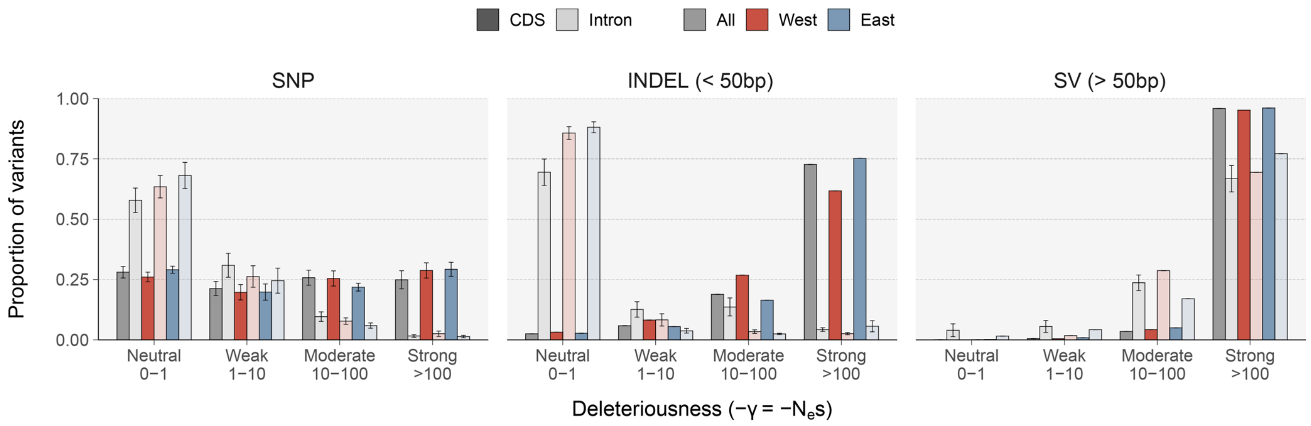


**Fig. S10.** Distribution of fitness effect (DFE) for the western and eastern populations, and for all samples, as inferred by the Anavar program.

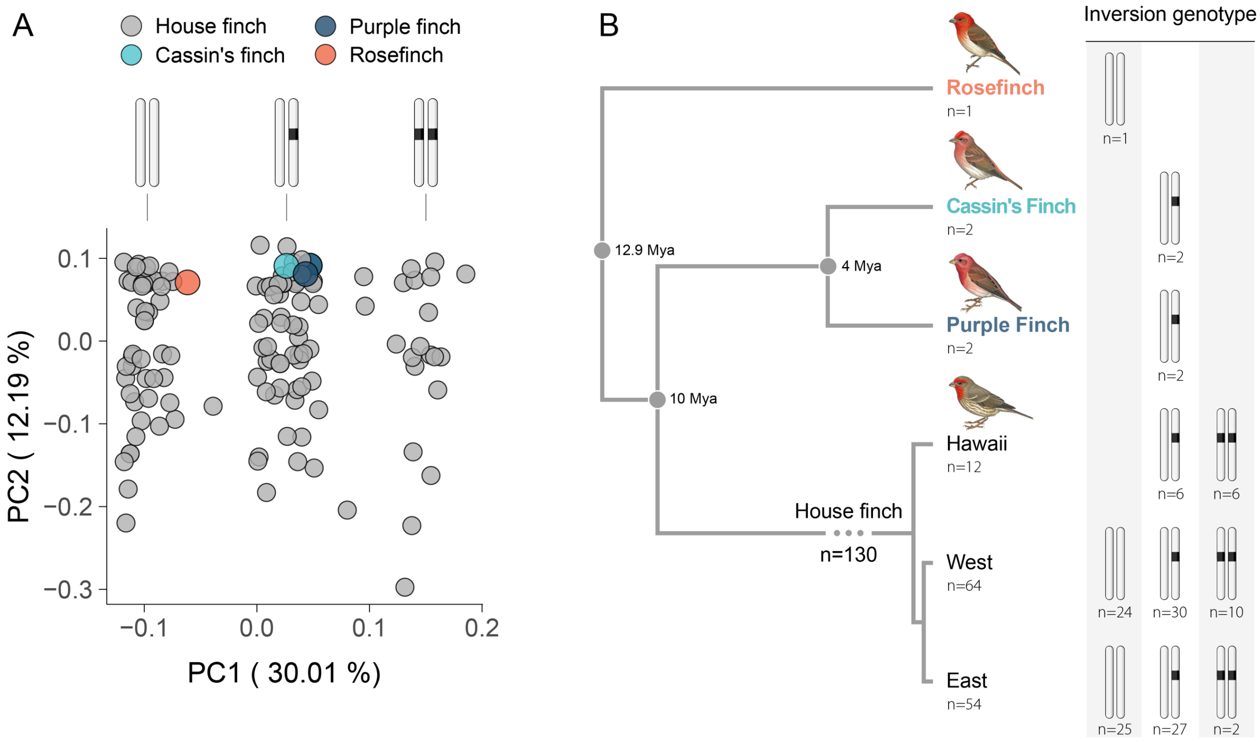


**Fig. S11.** Inversion haplotypes in the House Finch and outgroup samples in the study. (A) PCA based on SNPs within the inversion, indicating three inversion genotypes: homozygote, heterozygote and alternative homozygote. The homozygous inversion was determined as the ancestral (standard) haplotype based on the inversion genotype of the Rosefinch sample, following their phylogenetic relationships as shown in (B). (B) also demonstrates the inversion genotypes identified for the 135 samples used in the study, including 130 House Finches and five outgroup samples. The phylogenetic tree is reconstructed based on published studies (50, 57-60). Bird illustrations © Lynx Edicions.

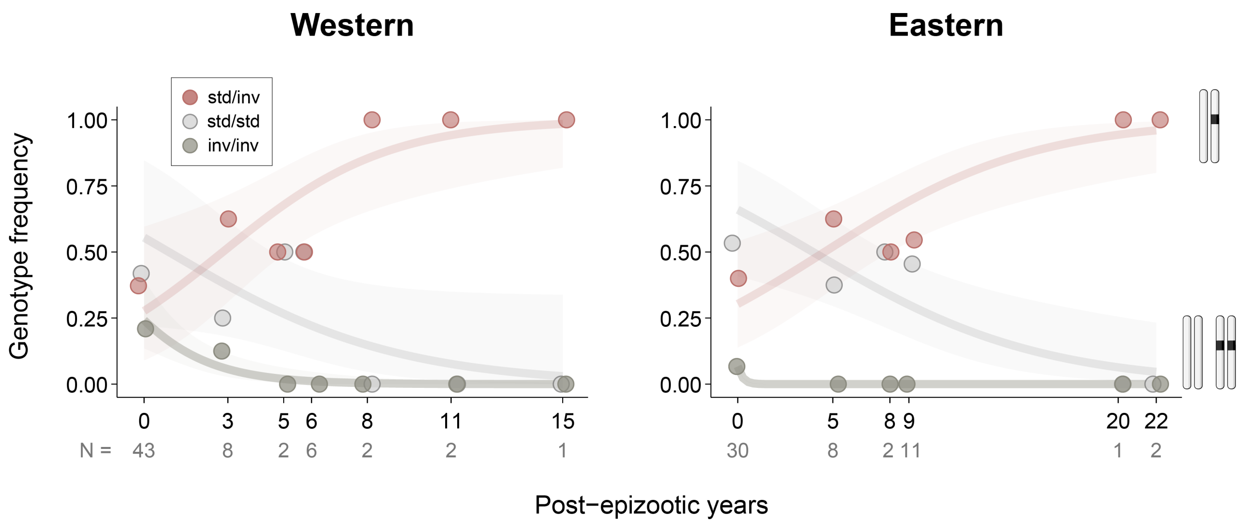


**Fig. S12.** Association between inversion genotype frequency and time since pathogen exposure in the western and eastern populations.

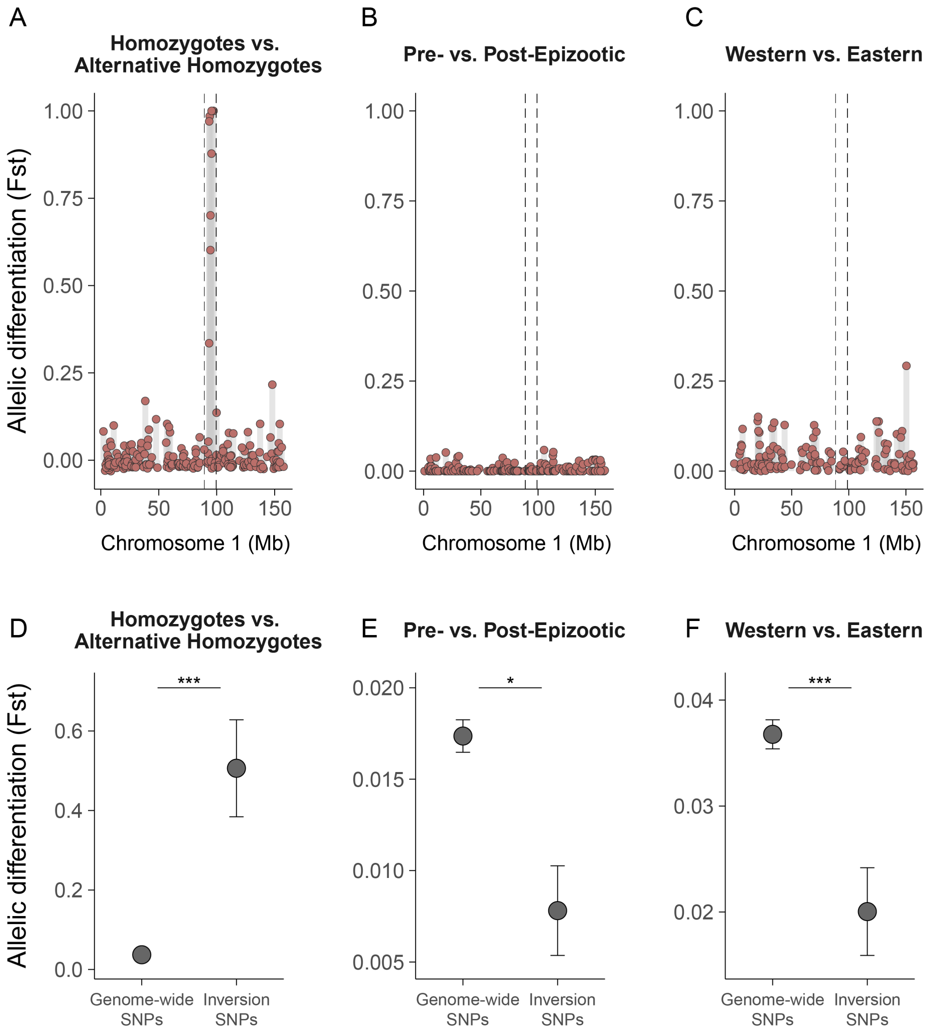


**Fig. S13.** Allele differentiation ($F_{ST}$; mean ± S.E.) between individuals with two different homozygous inversion (A, D), and between pre- and post-epizootic (B, E), and between western and eastern individuals (C, F). The inversion region is marked with dashed lines.

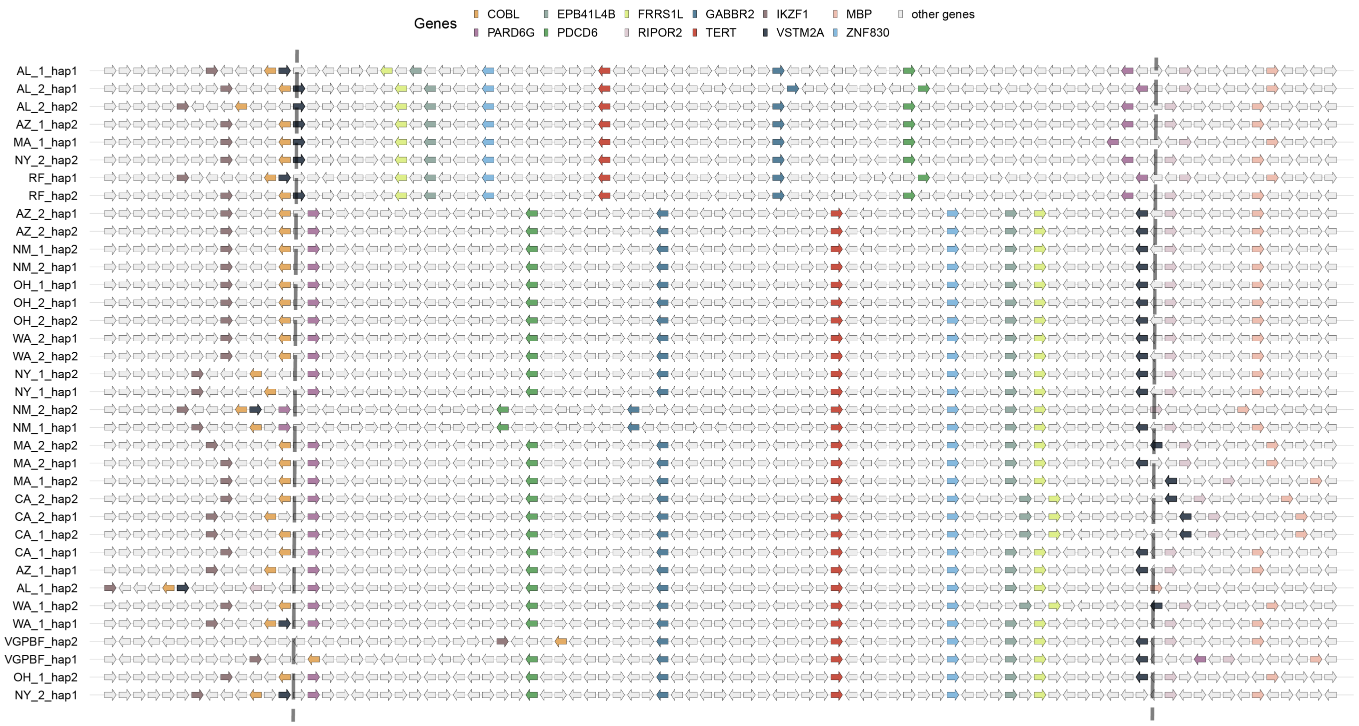


**Fig. S14.** Linear gene arrangements in the inversion and nearby breakpoints in all 32 haplotypes.

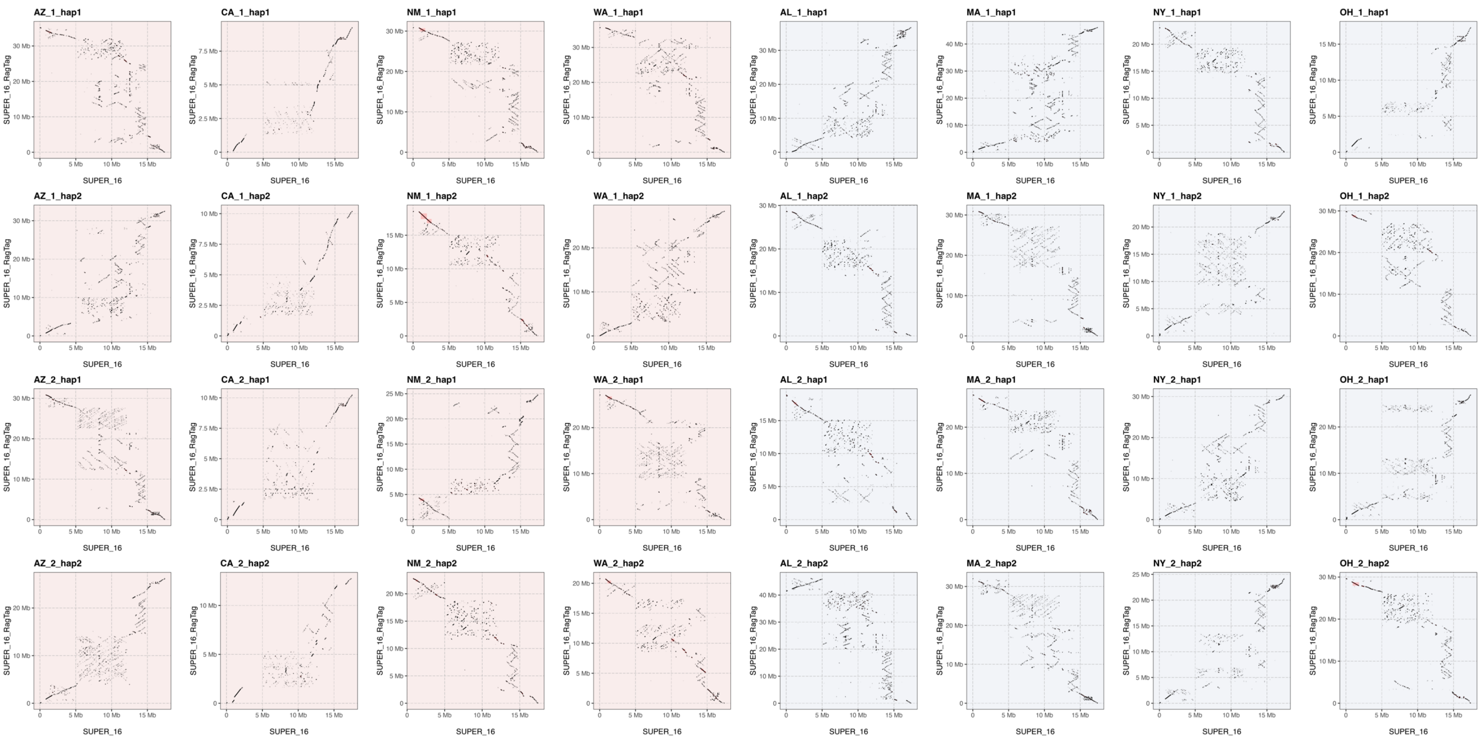


**Fig. S15.** Dot plots of chromosome 16 for each haplotype, displaying the alignment of each haplotype ('query', y-axis) with the VGP genome ('reference', x-axis). Chromosome 16 is composed of an extraordinarily large number of repeats.
